# Brain-wide Genome Editing via STEP-RNPs for Treatment of Angelman Syndrome

**DOI:** 10.1101/2025.11.13.684643

**Authors:** Xiaona Lu, Ying Xie, Youmei Bao, Jiang Yu, Zefeng Wang, Jiali Fan, Wendy C. Sheu, Gretchen Long, Zewei Tu, Jie Tong, Garret Manquen, Yichao Li, Varun Katta, Ryan Chow, Binfan Chen, Zhouqi Meng, Ying-hong Ma, Caihong Qiu, Sidi Chen, Shengdar Q. Tsai, Bony De Kumar, Caroline E. Hendry, Yong-hui Jiang, Jiangbing Zhou

**Affiliations:** Department of Genetics, Yale University School of Medicine, New Haven, CT, 06510, USA; Department of Neurosurgery, Yale University School of Medicine, New Haven, CT, 06510, USA; Department of Biomedical Engineering, Yale University, New Haven, CT, 06511, USA; Radiology & Biomedical Imaging, Yale University, New Haven, CT, 06510, USA; Department of Hematology, St Jude Children’s Research Hospital, Memphis, TN 38105, USA; Department of Medicine, Hospital of the University of Pennsylvania, Philadelphia, PA, USA; Yale Stem Cell Center, Yale University School of Medicine, New Haven, CT, 06510, USA; Department of Neuroscience, Yale University School of Medicine, New Haven, CT, 06510, USA; Department of Pediatrics, Yale University School of Medicine, New Haven, CT, 06510, USA; Department of Ob/Gyn and Reproductives Science, Yale University School of Medicine, New Haven, CT, 06510, USA; Wu-Tsai Institute, Yale University. 100 College Street, New Haven CT 06510

## Abstract

Brain-wide genome-editing remains a major hurdle for the treatment of neurogenetic disorders. Here, we report a non-viral, non-nanoparticle, chemical modification-based method, called Stimuli-Responsive Traceless Engineering Platform (STEP) that achieves highly efficient and brain-wide genome-editing of neurons in mice. Using cholesterol-based STEP as a lead, we show that a single administration of STEP-ribonucleoproteins (RNPs) results in functional genetic rescue with significant improvements across a battery of neurobehavioral domains in the Angelman syndrome (AS) mouse model. No significant off-target events or general toxicity effects are observed. STEP-RNPs are also highly efficient at editing human neurons and cortical brain organoids differentiated from AS patient-derived iPSCs. scRNA-seq analysis confirms functional genetic rescue via reactivation of *Ube3a/UBE3A* expression in human and mouse STEP-RNP-treated neuronal cells. Genome editing via STEP-RNPs has broad applications and the potential to treat many other neurogenetic disorders.

## Introduction

Genome editing has emerged as an exciting therapeutic strategy for genetic diseases^1–4^. Nevertheless, the safe and efficient delivery of genome-editing machinery, particularly to the brain in the context of neurogenetic diseases, remains a significant hurdle. Direct delivery of the CRISPR/Cas9 system as a ribonucleoprotein (RNP) complex, composed of a Cas9 protein and a single guide RNA (sgRNA), has emerged as an attractive approach for genome editing from a therapeutic perspective ^5–11^. However, current Cas9 RNP-based methods are not able to achieve widespread genome editing across the entire brain due to poor dispersion in brain tissues ^8,12–14^. While adeno-associated virus (AAV, ∼25 nm size)-mediated delivery of Cas9 DNA navigates some hurdles ^15–21^, their immunogenic nature, requirement of higher dose, suboptimal biodistribution with the currently utilized capsid in large animal brains and human applications, and the higher risk of off-target effects from persistent expression of Cas9 DNA, along with the potential risk of genomic integration and cancer development, present challenges for clinical application ^22,23^. In current clinical trials involving delivery of AAV-based therapies, either prophylactic or reactive immunosuppression protocols have been recommended for variable periods of time in individual trials, adding to the clinical challenge ^24–27^. Developments in nanoparticle-based delivery are promising; however, their considerable size, typically within the range of 40-300 nm, limits their dispersion within brain tissue where the extracellular space (ECS) is no wider than 35 nm ^11,28–31^. Delivery of RNPs has also been attempted through complexation with cationic peptides ^6,7^. Unfortunately, due to their high positive charge and limited stability, their application towards brain-wide editing via blood or cerebrospinal fluid (CSF) administration may also be limited ^14^. To overcome these limitations, we developed a highly efficient approach called stimuli-responsive traceless engineering platform (STEP) for the delivery of genome editing machinery in the form of RNP. STEP-RNP delivery is enabled through surface engineering of RNPs with heterofunctional chemicals of ∼2 kDa that are designed to interact with both the RNPs and target cells. The small size (12-14 nm) of STEP-RNPs is capable of achieving widespread brain distribution via CSF delivery. We reasoned that Angelman syndrome is an excellent model to test STEP-RNP genome editing.

Angelman syndrome (AS) is a severe neurodevelopmental disorder caused by maternal deficiency of the *UBE3A* gene in the chromosome 15q11.2-q13 region ^32,33^. The expression of the *UBE3A* gene is subject to brain-specific genomic imprinting ^34^. In the neurons of normal individuals, *UBE3A* is exclusively expressed from the maternal chromosome while paternal *UBE3A* expression is silenced by the paternally expressed non-coding transcript that is antisense to *UBE3A* (*UBE3A-ATS*) ^35,36^ though the exact molecular mechanism by which *UBE3A-ATS* silences *UBE3A* on the paternal chromosome is not well understood. Nonetheless, patients with AS typically have at least one structurally intact - albeit repressed - copy of *UBE3A* on the paternal chromosome. Disruption of *Ube3a-ATS* by modified antisense oligonucleotides (ASOs) ^37^ and viral vectors encoding CRISPR/Cas9 or Cas13 genome editing machinery has been shown to unsilence the expression of paternal *UBE3A,* as have small molecules and “artificial transcription factor” (ATF) approaches ^19,38–41^. All of these strategies, some of which are transient for the reactivation of *UBE3A*, have shown positive signs of clinical efficacy in the AS mouse models. Similarly, strong positive signals have been observed in multiple behavioral domains from three independent phase 1/2a ASO clinical trials ^42,43^, with Phase 3 trials launched by Ultragenyx and Ionis in late 2024. However, ASOs have only a transient effect *in vivo* due to their short half-life in brains and mRNA targeting mechanism, which necessitates frequent intrathecal (IT) administration. Similarly, CRISPR RNA editing strategies via Cas13, small molecule, and ATF, if successful in clinical trials, are also expected to require frequent repeat administration ^44^. Our non-viral and non-nanoparticle-based genome editing STEP-RNP delivery platform is poised to overcome both these limitations by facilitating permanent genome editing to disrupt the *Ube3a-ATS* brain-wide, whilst simultaneously minimizing the safety risks associated with virus-mediated delivery. Since our STEP-RNP technology can accommodate any sgRNA, it offers a novel genome editing platform for many neurogenetic and also non-neurogenetic disorders.

## Results

### Synthesis and characterization of STEP-RNPs

To achieve efficient delivery of RNPs, we developed a novel chemical engineering approach that does not rely on nanocarriers or peptides but instead utilizes STEP molecules. STEP molecules comprise two key components: the cell-penetration module, which ends with a small molecule, R1, that facilitates entry into the cell, and the protein-interaction module, which ends with a chemical structure, R2, selected for interaction with protein surfaces. These two modules can be chemically joined together, forming a complete STEP chemical. This design enables STEP molecules to interact with both the target cells (via R1) and the RNPs (via R2), thereby facilitating the cytosolic delivery of genome-editing machinery (**Fig. 1a**).

**Fig 1.**
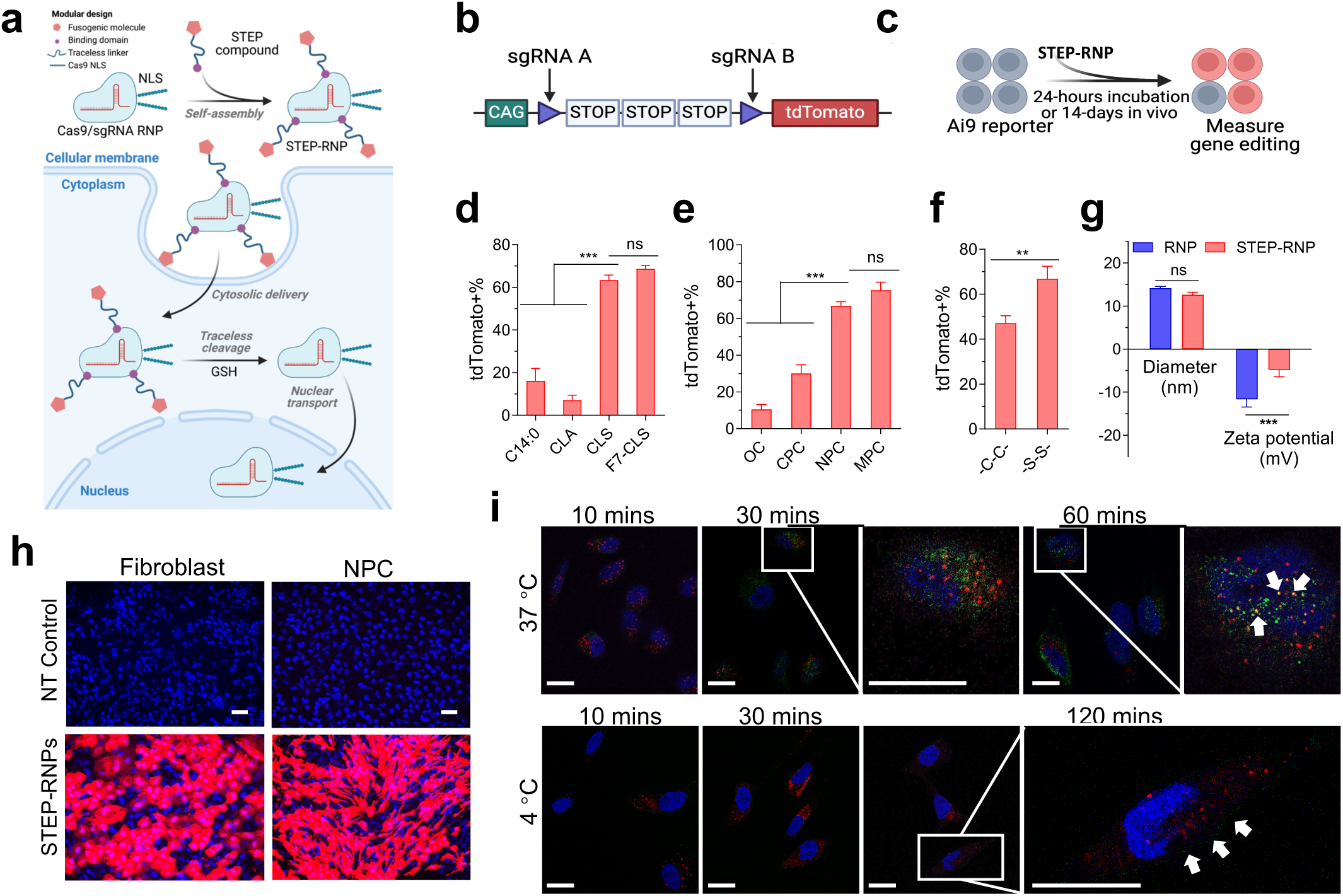
Design and characterization of the STEP-RNP platform. **a.** STEP binds to pre-assembled RNP and is taken up by the cell via endocytosis. Elevated intracellular glutathione (GSH) levels trigger cleavage of the disulfide linkage, resulting in STEP detachment. Nuclear localization signals (NLS, green) facilitate subsequent nuclear import of the RNP complex. **b.** Diagram of the *Ai9* reporter construct containing three STOP cassettes flanked by loxP sites upstream of the *tdTomato* reporter. Two guide RNAs (sgRNA-A and sgRNA-B) target regions immediately upstream and downstream of the STOP cassette; successful editing at both sites excises the STOP sequence, enabling *tdTomato* expression. **c.** Schematic of STEP-RNP–mediated editing in *Ai9*-derived reporter cell models. **d.** Screening of R1 modules for genome editing efficiency in *Ai9* reporter cells (Cas9, 10 µg/mL). Cholesterol (CLS) and F7-Cholesterol (F7-CLS) exhibited significantly higher efficiency compared to 14 other R1 candidates (\*\*\**p* < 0.001; *n* = 3; see also Extended Data Fig. 1a,b). **e.** Evaluation of R2 binding modules for genome editing delivery in *Ai9* reporter cells (Cas9, 10 µg/mL). NPC (4-Nitrophenyl Carbonate) and MPC (Methoxyphenyl Carbonate) showed significantly enhanced editing efficiency relative to six other binding modules (\*\**p* < 0.01; *n* = 3; see also Extended Data Fig. 1c,d). **f.** Comparison of cleavable (-S-S-) versus non-cleavable (-C-C-) linkage pairs revealed significantly higher editing efficiency for the cleavable configuration (\*\**p* < 0.01; *n* = 3; see also Extended Data Fig. 1f-i). **g.** Dynamic light scattering analysis of STEP-RNP complexes showing an average hydrodynamic diameter of 13 nm and a negative surface zeta potential. **h.** Representative fluorescence images of embryonic fibroblasts and neural progenitor cells (NPCs) derived from *Ai9* reporter mice treated with STEP-RNPs carrying either scrambled gRNA (STEP-RNP-sgRNA-NC, top) or target-specific gRNAs (STEP-RNP-sgRNA-AB, bottom) (*n* = 20; scale bar = 50 µm). **i.** STEP-RNP enters cells via endocytosis. HeLa cells incubated with STEP-RNP-GFP (10 µg/mL) were stained for DAPI and the endosomal marker EEA1. STEP-RNP-GFP colocalized with EEA1 at 37 °C (arrows), but not at 4 °C, indicating temperature-dependent endocytosis. Representative images from three independent experiments (scale bar = 20 µm).

To identify an optimized STEP molecule for RNP delivery, a library of molecules was synthesized by reacting an array of different R1 modules selected to facilitate cell-penetration (**Extended Data 1a)**, with a suite of different R2 protein-interaction modules (**Extended Data 1c**). To screen for the most efficient genome-editing STEP formulation, we assembled RNPs using Cas9 with sgRNA A and B targeting the loxP sites of the stop cassette within the Ai9 transgene, when removed, activates expression of tdTomato ^45,46^ **(Fig. 1b,c**). STEP-RNPs were incubated with mouse embryonic fibroblasts (MEFs) derived from Ai9 reporter mice for 24 hours and editing efficiency was assessed based on tdTomato expression 48 hours later. Among all the tested R1 molecules, cholesterol and its analog, exhibited the highest efficiency (**Fig. 1d & Extended Data 1b**). Cholesterol was chosen due to its endogenous presence in humans, which we reasoned would pose a lower safety risk. Among all the R2 molecules, 4-nitrophenyl carbonate (NPC) demonstrated the greatest efficiency and was therefore selected (**Fig. 1e & Extended Data 1d**). We compared a group of linker molecules and found that STEP molecules with a cleavable disulfide bond (-S-S-) within the protein-interaction module showed superior delivery efficiency compared to non-cleavable carbon-carbon bond (-C-C-). (**Fig.1f** & **Extended Data. 2a,b)**. This enhancement is likely due to the cleavage of the disulfide bond by glutathione (GSH), which is abundant in the cytoplasm, allowing for traceless release of the RNP payload. The efficiency of STEP-mediated delivery is also influenced by the length of the STEP molecule (**Extended Data 2c,d**).

We sought to optimize the formulation to maximize genome-editing efficiency. The efficiency of STEP for delivering RNPs was correlated with the mole ratio of STEP molecule to RNP, with an optimal ratio of 20 for editing Ai9 reporter MEFs (**Extended Data 2e**). Ratios exceeding 20 did not further enhance editing efficiency. Cells remained fully viable at a STEP-RNP ratio of 20 and Cas9 concentration up to 10 µg/mL for Ai9 MEFs and 15 ug/mL for Ai9 neural progenitor cells (NPCs) (**Extended Data 2f,g**). Complexing STEP at a ratio of 20 reduced the surface charge without significantly increasing the size of RNPs. The final STEP-RNPs is approximately 13 nm in diameter and carries a negative charge (**Fig 1g** & **Extended Data 2h,i**). Treatment with RNPs at a concentration of 10 µg per ml of culture volume achieved 80.9±0.7% editing in Ai9 MEFs and 73.0±5.2% in NPCs (**Fig. 1h**& **Extended Data 2j**). The efficiency of STEP-RNPs also depends on the Cas9 variant used, as STEP-RNPs assembled with G5, showed significantly higher efficiency than V3-based STEP-RNPs (**Extended Data 2k**). The STEP-RNP complexes were highly stable and maintained their editing efficiency in artificial cerebral spinal fluid (ACSF) and serum-containing medium at 37°C for over 24 hours or at -20°C for over 4 months (**Extended Data 2l)**. Using Cas9 fused with GFP as a payload, we observed that most GFP-RNPs successfully penetrated the nuclei within 60 minutes in both Hela cells and NPCs (**Fig. 1h & Extended Data 3**). However, incubation at 4°C, a temperature at which endocytosis is negligible (*40*), or co-treatment with Pitstop 2, a potent inhibitor of Clathrin-dependent endocytosis, significantly reduced cell penetration of STEP-RNP-GFPs (**Fig. 1i &Extended Data 2m)**. Based on these data, we hypothesize that STEP uptake is mediated via the endocytosis pathway followed by entrance into the nucleus, however a more detailed study of cellular uptake remains to be investigated (**Fig. 1i & Extended Data 2m,3**).

### Brain-wide distribution of STEP-RNPs in the mouse via systemic delivery

To assess the in vivo editing potential of STEP-RNPs, we performed a local intracranial administration of STEP-RNPs through convection-enhanced delivery (CED) to the striatum of Ai9 reporter mice (**Fig. 2a**). The striatum was chosen due to its optimal feasibility and capacity for higher injection volumes. As expected for CED, dispersion was local around the injection site, with extensive dispersion and efficient editing observed in both NeuN positive neurons and GFAP positive astrocytes (**Fig. 2b-e**).

**Fig 2.**
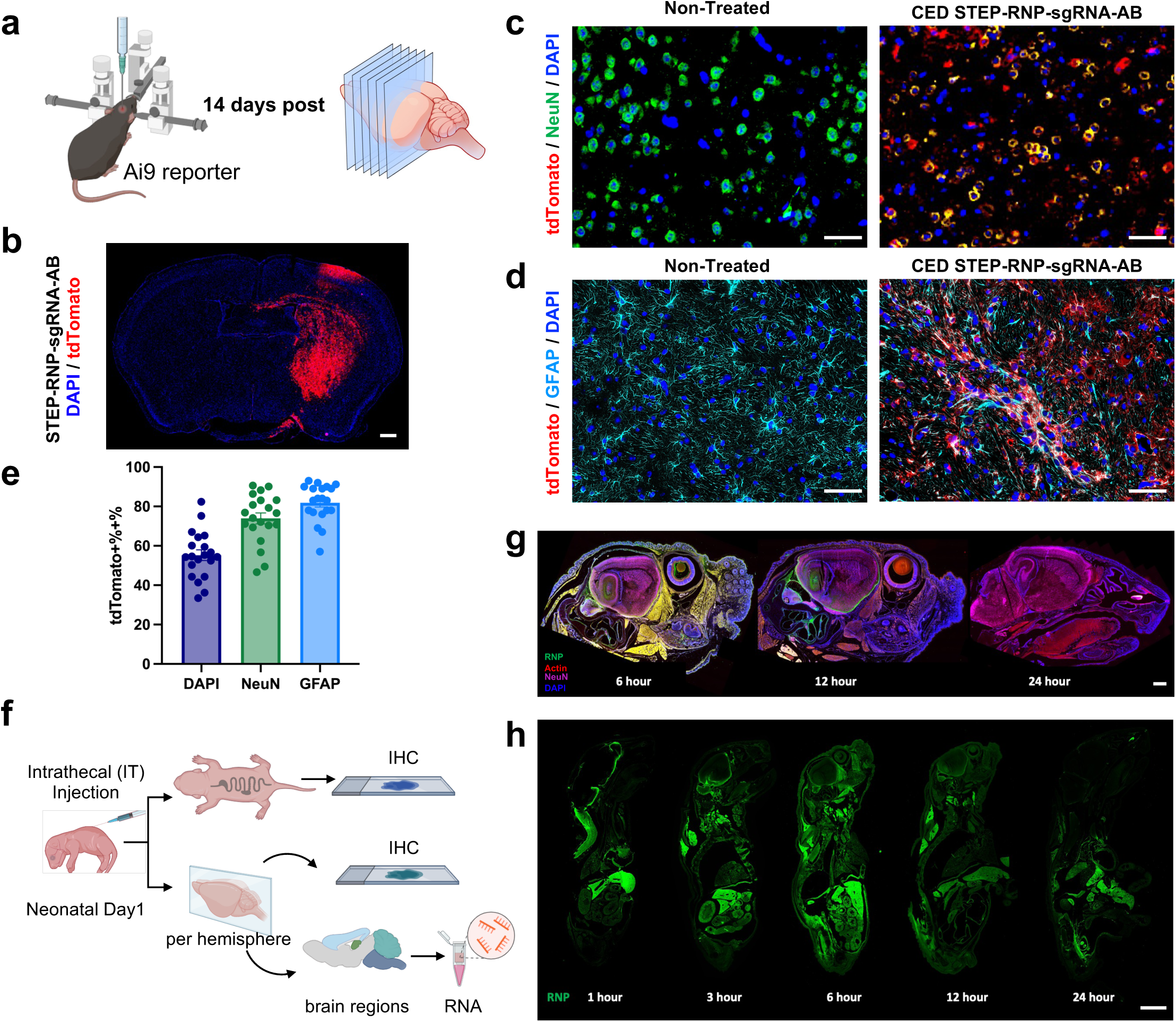
STEP-RNP mediated local editing in striatum and biodistribution in mouse brain. **a.** Schematic of the experimental design illustrating convection-enhanced delivery (CED) of STEP-RNP into the striatum to test in vivo gene-editing potential. **b.** Efficient gene editing within the injected striatal region visualized by tdTomato reporter activation (scale bar = 500 µm). **c-d.** Co-immunostaining of tdTomato (red) with the neuronal marker NeuN (green; c) and the astrocytic marker GFAP (blue; d) demonstrating neuronal and glial transduction (*n* = 20; scale bar = 50 µm). **e.** Quantification of total gene-edited cells (DAPI+), neurons (NeuN+), and astrocytes (GFAP+) following CED-mediated delivery of STEP-RNP-sgRNA-AB in *Ai9* reporter mice. **f.** Schematic of the biodistribution study of STEP-RNP following intrathecal (IT) administration. **g-h.** Representative images showing STEP-RNP localization in the neonatal mouse brain (g) and whole body (h) at 1, 3, 6, 12, and 24 hours post-IT injection (10 µg/µL, 10 µL/mouse). STEP-RNP was detected by immunostaining with anti-Cas9 antibody and observed in multiple brain regions, including the midbrain, hypothalamus, and pons-medulla, within 1 hour of injection. Signal intensity peaked at 6 hours and was undetectable in the brain by 24 hours (*n* = 12 per time point; scale bars: 500 µm in g, 5 mm in h).

To assess the in vivo biodistribution of STEP-RNPs in the brain, we administered STEP-RNPs to postnatal day 1 (P1) pups via IT injection (**Fig. 2f**). Immunostaining using an anti-Cas9 antibody at various time-points post-injection showed that STEP-RNPs achieved widespread distribution in the brain and also circulated to peripheral tissues (**Fig. 2g,h**). Biodistribution peaked in the brain and non-brain tissues around 6-12 hours post-administration and became nearly undetectable in the brain and significantly reduced in non-brain tissues by 24 hours (**Fig. 2g,h**). We also tested intravenous (IV) injection of STEP-RNPs and observed wide and transient distribution in the brain but with a higher accumulation in the liver compared to IT route (**Extended Data 4a-c**).

We then assessed biodistribution and editing efficiency together using an AS-specific mouse model containing a *Ube3a-YFP* fusion construct that reliably reports de-repression and activation of the paternal *Ube3a* allele in neurons (*Ube3a^m+/p-YF^*^P^) (JAX stock #017765) ^47^ (**Fig.3a,b**). The *Ube3a* gene has several alternative promoters and alternative splicing that drive ubiquitous expression of different mRNA isoforms in mouse brain that are subjected to genomic imprinting in neurons by paternally expressed *Ube3a-ATS* ^48^*. Ube3a* is exclusively expressed from the maternal allele in neurons but biallelically in glia cells^48,49^. Thus, the Ube3a-YFP reporter line provides a highly specific tool to assess editing efficiency in neurons and examine paternal specific reactivation of *Ube3a* using hybrid PCR primers anchored to *Ube3a* and the *YFP* coding region^37,40,41^.

To identify an appropriate sgRNA, we first redesigned and screened sgRNAs targeting *Ube3a-ATS* between *Ube3a* and *Snord115* based on previously published sequences ^41,50^ (**Extended Data 5a**). Of the different sgRNAs administered, sgRNA33 demonstrated superior efficiency in reducing the *Ube3a-ATS* as measured by primer pairs at multiple loci in brain 30 days post-delivery (**Fig. 3c**).

**Fig 3.**
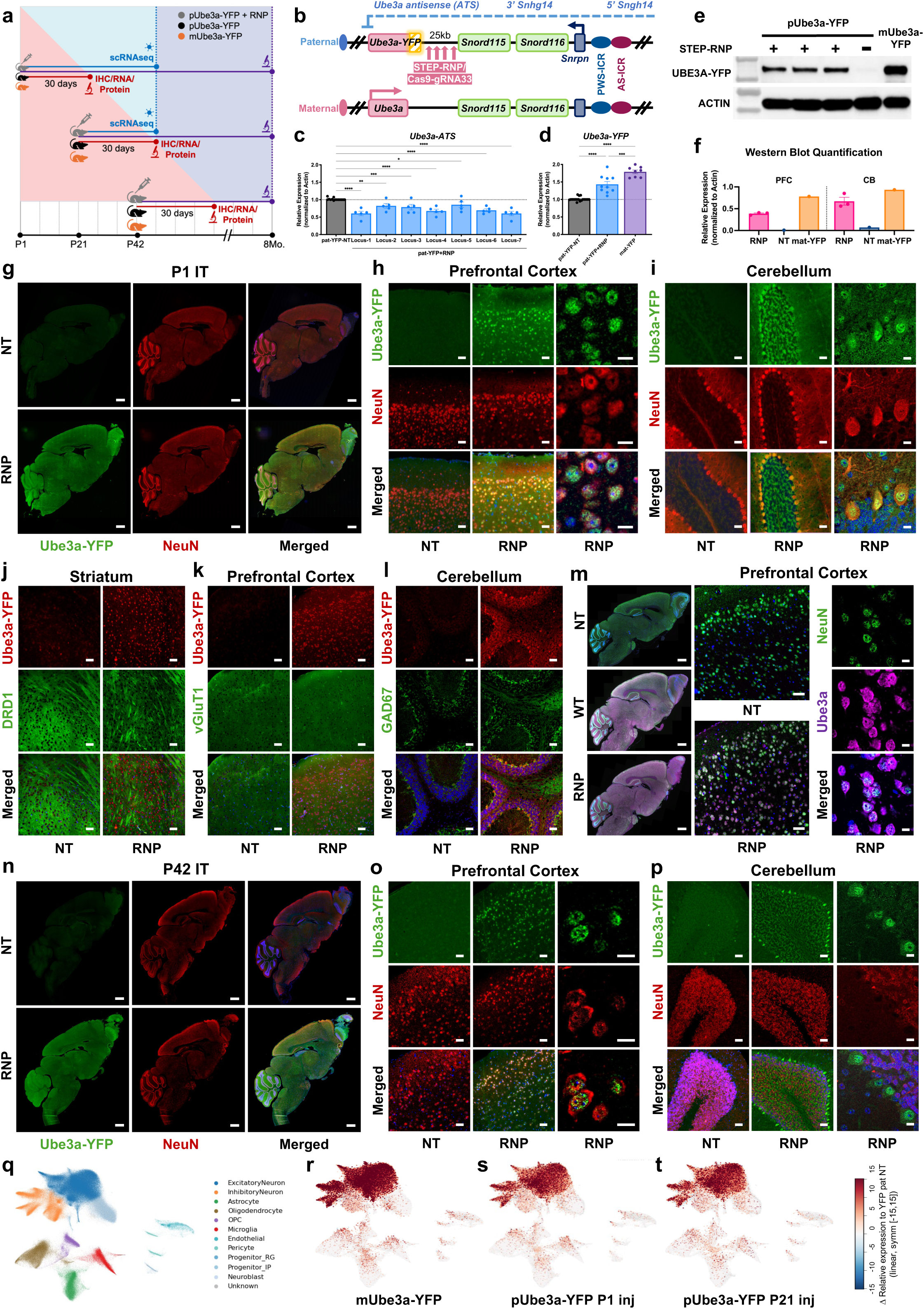
STEP-RNP mediates brain-wide editing and reactivation of paternal *Ube3a-YFP* in Angelman syndrome reporter mice. **a.** Schematic of experimental design used to evaluate editing efficiency in Ube3a-YFP reporter mice. **b.** Illustration of the paternal *Ube3a-YFP* locus showing the STEP-RNP editing sites and imprinting regulation involving *Snhg14* and *Ube3a-ATS* across the region spanning the imprinting center to *Ube3a*. **c.** Reduction of *Ube3a-ATS* level assessed by qRT-PCR 30 days after P1 IT administration of STEP-RNP. **d.** Reactivation of *Ube3a-YFP* mRNA detected by qRT-PCR 30 days after P1 IT administration of STEP-RNP. **e-f.** Reactivation of UBE3A-YFP protein confirmed by immunoblot 90 days after P1 STEP-RNP IT delivery. Maternal UBE3A-YFP served as the positive control, and paternal UBE3A-YFP without treatment as the negative control. Quantification in f. **g.** Representative fluorescence images showing brain-wide reactivation of the paternal UBE3A-YFP 30 days after IT injection of STEP-RNP (G5 + sgRNA33, 10ug/ul, 10uL/mouse) into P1 *Ube3a^m+/p-YF^*^P^ mice (scale bar = 1 mm). **h-i.** Reactivation of paternal UBE3A-YFP in the prefrontal cortex (PFC) and cerebellum 30 days after P1 IT injection (scale bar = 50 µm in 1^st^ & 2^nd^ column, scale bar = 10 µm in 3^rd^ column). **j-l.** Colocalization of reactivated paternal UBE3A-YFP with neuronal subtype markers, including dopaminergic neurons (DRD1), glutamatergic neurons (vGLUT1), and GABAergic neurons (GAD67), across the striatum, PFC, and cerebellum (*n* = 6; scale bar = 50 µm). **m.** Sustained reactivation of paternal UBE3A-YFP in the PFC 8 months after P1 STEP-RNP injection (scale bar: 1 mm in 1^st^ column, 50 µm in 2^nd^ column, 10 µm in 3^rd^ column). **n.** Representative images showing brain-wide reactivation of paternal UBE3A-YFP 30 days after IT injection (10 µg/µL, 10 µL/mouse) of STEP-RNP-AS into adult (P42) *Ube3a^m+/p-YF^*^P^ mice (scale bar = 1 mm). **o-p.** Reactivation of paternal UBE3A-YFP in PFC and cerebellum 30 days after P42 IT injection (scale bar = 50 µm in 1^st^ & 2^nd^ column, scale bar = 10 µm in 3^rd^ column). **q.** Single-nucleus RNA-seq profile of *Ube3a* expression across brain cell types. **r-t**. Expression of maternal *Ube3a*-*YFP* (r) and reactivation of paternal *Ube3a*-*YFP* in the cortex of P1 (s) and P21 (t) mice following IT delivery of STEP-RNP. Expression values represent single-nucleus *Ube3a*-*YFP* levels after subtracting the median expression of paternal *Ube3a*-*YFP* in non-treated mice.

We delivered STEP-RNPs with various sgRNAs to P1 pups via IT or intracerebroventricular (ICV) injections. We found that ICV at P1 yielded comparable results to IT (**Extended Data 5b**), which were also more efficacious than IV (**Extended Data 4e-h**). As such, IT injection was selected for STEP-RNP delivery in subsequent experiments due to the relative ease of administration ^51^ and lower risk of damage caused by injection of the needle into the brain for clinical efficiency studies.

Further analysis of the effect of STEP-RNP-delivered sgRNA33 demonstrated effective reactivation of the paternal *Ube3a-YFP* gene without significantly affecting the expression of surrounding genes, particularly *Snord116*, whose reduction from the paternal chromosome could potentially implicate Prader-Willi syndrome ^52,53^ (**Extended Data 5c**). Quantitative RT-PCR analysis showed significant reactivation of *Ube3a*-YFP in the brain 30 days post-injection (**Fig. 3d**). Immunostaining analysis at the same timepoint revealed widespread detection of UBE3A-YFP in a high percentage in NeuN+ neurons (77.8±3.1%) in the neocortex and in calbindin+ Purkinje cell (80.5±5.8%) in the cerebellum (**Fig 3g-i**), further supportive of successful editing and reactivation of UBE3A-YFP. We also observed robust reactivation and expression of UBE3A-YFP in DRD1^+^, vGluT1^+^, and GAD67^+^ neurons in the striatum, prefrontal cortex and cerebellum respectively (**Fig. 3j-l**). Delivery of the STEP-RNPs to mice at 42 days of age achieved similar results (**Fig. 3n-p**). At 90 days post-administration, UBE3A reactivation reached 49.5±2.2% of the maternal expression level in the prefrontal cortex and 71.8±9.7% in the cerebellum as assessed by quantitative immunoblot (**Fig. 3e,f**). Our final observation point was 8 months post P1 IT STEP-RNP delivery, at which time we continued to observe sustained reactivation and expression of UBE3A-YFP (**Fig. 3m**). We were not able to assess the editing efficacy in non-neuronal cells using Ube3a-YFP reporter line due to the biallelic expression of *Ube3a* in these cells^48,54^. Consistent with these results, we also observed brain-wide editing of STEP-RNPs in Ai9 tdTomato reporter mice using both Cas9 variants of G5 and G221 (**Extended Data 4d**).

To further support evidence of successful gene editing, we performed single-nucleus RNA-seq of cerebral cortex in *Ube3a^m+/p-YF^*^P^ treated (or not) with STEP-RNPs. We reasoned that if gene editing is successful, *Ube3a-YFP* expression in neurons should be reactivated to levels approaching the maternal allele. Upon IT administration of STEP-RNPs either at P1 (**Fig. 3s**) or P21 (**Fig. 3t**), we observed strong expression of *Ube3a-YFP* specifically in excitory and inhibitory neurons, suggesting successful editing and reactivation of *Ube3a* in these mice compared to *Ube3a^p+/m-YF^*^P^ (m-Ube3a-YFP) control (**Fig.3r**). These results demonstrate that the STEP platform achieves highly efficient and long-lasting editing of neurons in mouse brains.

### STEP-RNPs ameliorate neurobehavioral impairments in an AS mouse model

Having demonstrated the potential for STEP-RNP-mediated gene editing in the brain in vivo, we next wanted to assess whether administration of STEP-RNPs could rescue neurobehavioral impairments in AS, a severe neurodevelopmental disorder with well-defined impairments, including seizures. To assess the optimal dose for clinical efficacy studies, we performed a dosage response study using the amount of Cas9 protein for the dosage calculation via IT administration at P21 (**Extended Data 5d-i**). Molecular analysis revealed the reactivation of *Ube3a-YFP* saturated at the dose of 75 ug (50%) of Cas9 **(Extended Date 5d**). We also found that 150 ug (100%) of Cas9 is the minimal effective dose for the majority of behavioral rescues (**Extended Data 5e-i**).

We next investigated whether treatment with STEP-RNPs at a 150 ug dose could mitigate the neurobehavioral impairments observed in AS mice with maternal *Ube3a* deficiency on a pure C57BL6/J background (*Ube3a^m-/p+^*) (JAX stock #016590) ^55^. For these experiments, a single dose of STEP-RNPs formulated with either V3 Cas9 or G5 Cas9 variant (Methods) was administered to mice in two separate cohorts: neonatal (P1) and weaning or adult ages (P21/P42) (**Fig. 4a**). We found that the treatments reduced expression of *Ube3a-ATS* (**Fig. 4b**) and increased the expression of *Ube3a* (**Fig. 4c**). A battery of behavioral tests (described below) was conducted at specific time points to evaluate different functional behavioral domains including cognitive function across the treated group, non-treated (NT) group, control group treated with STEP-RNPs containing scrambled sgRNA and WT mice (**Fig. 4d**).

**Fig. 4.**
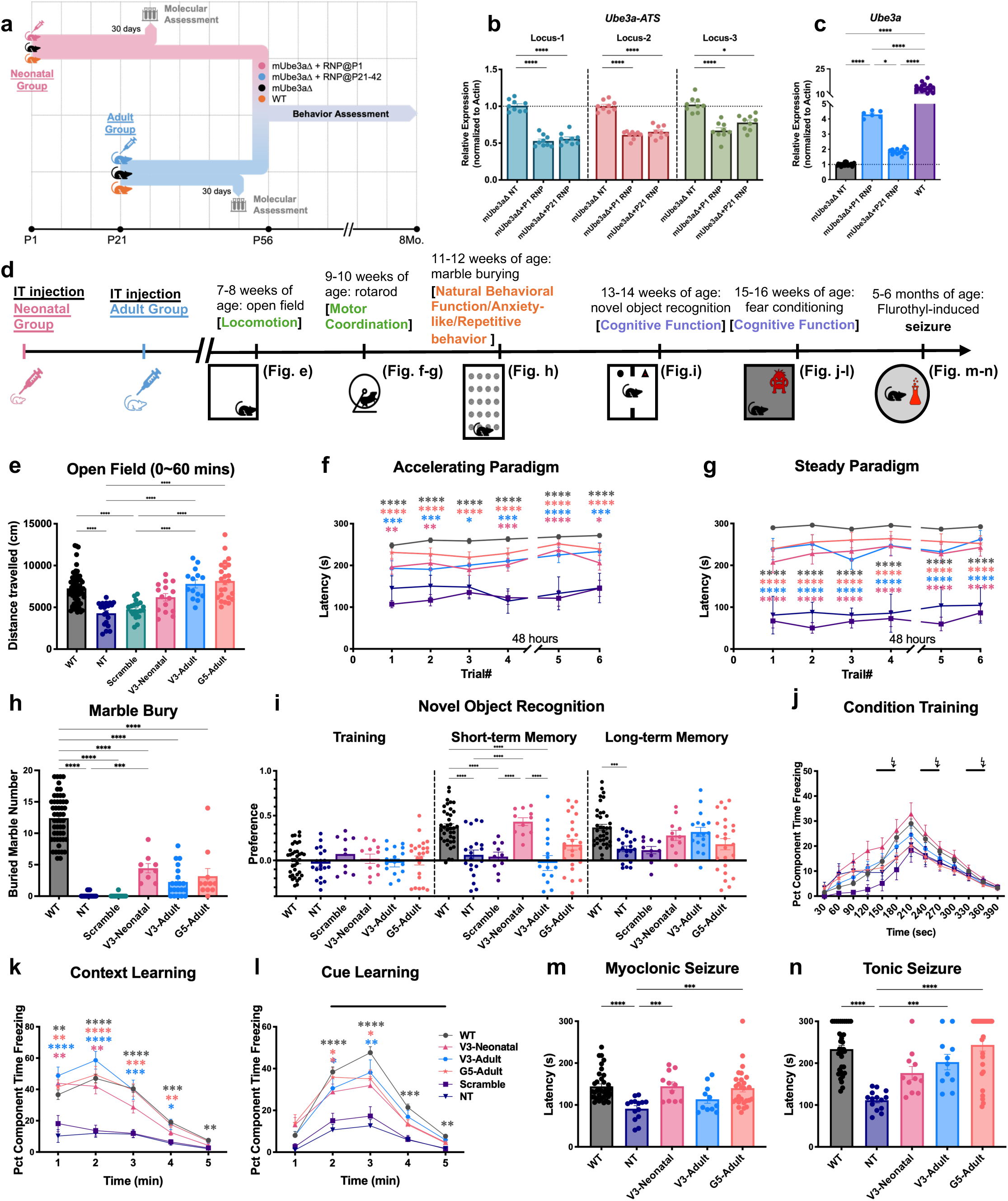
STEP-RNP editing restores *Ube3a* expression and rescues behavioral phenotypes in the Angelman syndrome (AS) mouse model (*Ube3a^m-/p+^*, mUbe3aι1) **a.** Schematic of the experimental design evaluating gene editing and therapeutic efficacy of STEP-RNP in the AS maternal *Ube3a* deficiency mouse model. **b.** Reduction of *Ube3a-ATS* levels in the prefrontal cortex (PFC) of *Ube3a^m-/p+^* mice 30 days after postnatal day 1 (P1) intrathecal (IT) administration of STEP-RNP, assessed by qRT-PCR (*n* = 4 per group; \**p*<0.05; ** *p*<0.01). **c.** Reactivation of *Ube3a* mRNA in the brains of *Ube3a^m-/p+^* mice 30 days after STEP-RNP P1 IT administration, measured by qRT-PCR (*n* = 4 per group; \**p*<0.05; ** *p*<0.01). **d.** Schematic overview of behavioral assessments designed to evaluate the functional efficacy of STEP-RNP in the AS mouse model (*Ube3a^m-/p+^*). Neonatal (P1, pink) and adult (P21/P42) cohorts received a single STEP-RNP injection prior to behavioral testing. WT, wild type; NT, no treatment. **e. Open-field test:** STEP-RNP treatment (V3 Cas9 or G5 Cas9) significantly improved locomotor activity compared to NT and scrambled gRNA controls. **f-g. Rotarod test:** STEP-RNP-treated AS mice exhibited significantly enhanced motor coordination under both accelerating (h) and steady-speed (i) conditions relative to untreated and scrambled controls. **h. Marble burying test:** STEP-RNP treatment partially restored marble-burying behavior compared to NT and scrambled groups. **i. Novel object recognition:** STEP-RNP treatment significantly improved short-term object memory, with complete rescue in P1-treated AS mice, and enhanced long-term memory performance across all treated groups. **j-l. Fear conditioning:** STEP-RNP-treated AS mice showed significantly improved contextual and cue learning compared to NT and scrambled controls at both age-groups. **m-n. Flurothyl-induced seizure test:** STEP-RNP treatment increased seizure latency, indicating reduced susceptibility to both myoclonic and tonic seizures. Notably, both neonatal and adult STEP-RNP-treated AS mice achieved full rescue of latency to wild-type levels. (\**p*<0.05, \*\**p*<0.01, \*\*\**p*<0.001).

Seizures are one of the most challenging clinical issues in patients with AS and can be a primary cause of mortality ^33,56,57^. We tested seizure susceptibility of AS mice at 5-6 months of age following STEP-RNP delivery at neonatal or adult stages, respectively. Untreated AS mice exhibit increased susceptibility to seizure induction ^55,58,59^; in contrast, administration of STEP-RNPs, regardless of the age at which it is administered, was effective in suppressing both myoclonic and tonic seizures (**Fig. 4m,n**). The administration of STEP-RNPs also ameliorated cognitive deficits in AS mice, as evidenced by improved performance in tests of short-term and long-term memory (**Fig. 4i**). Mice treated in the neonatal stage exhibited complete recovery in short-term memory assessments, whereas those treated in adulthood showed partial improvement. Furthermore, all groups, regardless of treatment timing, demonstrated moderate enhancements in long-term memory performance. Assessments of fear conditioning encompassing both contextual and cued paradigms revealed that STEP-RNP treatment nearly fully restored contextual learning in AS mice and significantly ameliorated cue learning (**Fig. 4j-l**).

AS patients exhibit anxiety, motor impairment and incoordination ^60^. In AS mice, we found that STEP-RNP treatment improved motor function and anxiety-like behavior in all tests. Treated AS mice exhibited an increased total distance traveled during a 1-hour open field test in both neonatal and adult groups (**Fig. 4e**). Improvements in motor coordination were evident from the prolonged latency to fall in both accelerating and steady-speed in repeat rotarod tests (**Fig. 4f-g**). Notably, with both V3 Cas9 and G5 Cas9, mice treated at adulthood significantly improved motor coordination. Additionally, STEP-RNP treatment resulted in partial correction of marble burying behavior (**Fig. 4h**), reflecting an improvement in proprioceptive function and anxiety-like behavior in AS mice. Together, these data provide strong evidence that STEP-RNPs are effective for the long-term amelioration of neurobehavioral deficits observed in AS model mice, including seizure susceptibility.

### Efficient human genome editing via STEP-RNPs in AS iPSC-derived neurons and organoids

The imprinting mechanism of *UBE3A* is highly conserved between humans and rodents, but the sequences and genomic structure of *UBE3A-ATS* are not ^61^. The identification of highly efficient human-specific sgRNAs (hsgRNAs) therefore requires a human-specific neuronal cell system in which to screen and optimize candidates. To this end, we engineered a *UBE3A-EGFP* fusion gene into the endogenous *UBE3A* paternal allele of an induced pluripotent stem cell (iPSC) line derived from an AS patient with a large maternal deletion of 15q11.2-q13. The EGFP was fused in-frame to the C-terminus of the paternal *UBE3A* gene before the STOP codon (**Fig. 5a & Extended Data 6a,b**). We then differentiated these cells into neurons (hereafter called AS-iPSC-iNs) following the NGN2 overexpression protocol ^62^ .

**Fig 5.**
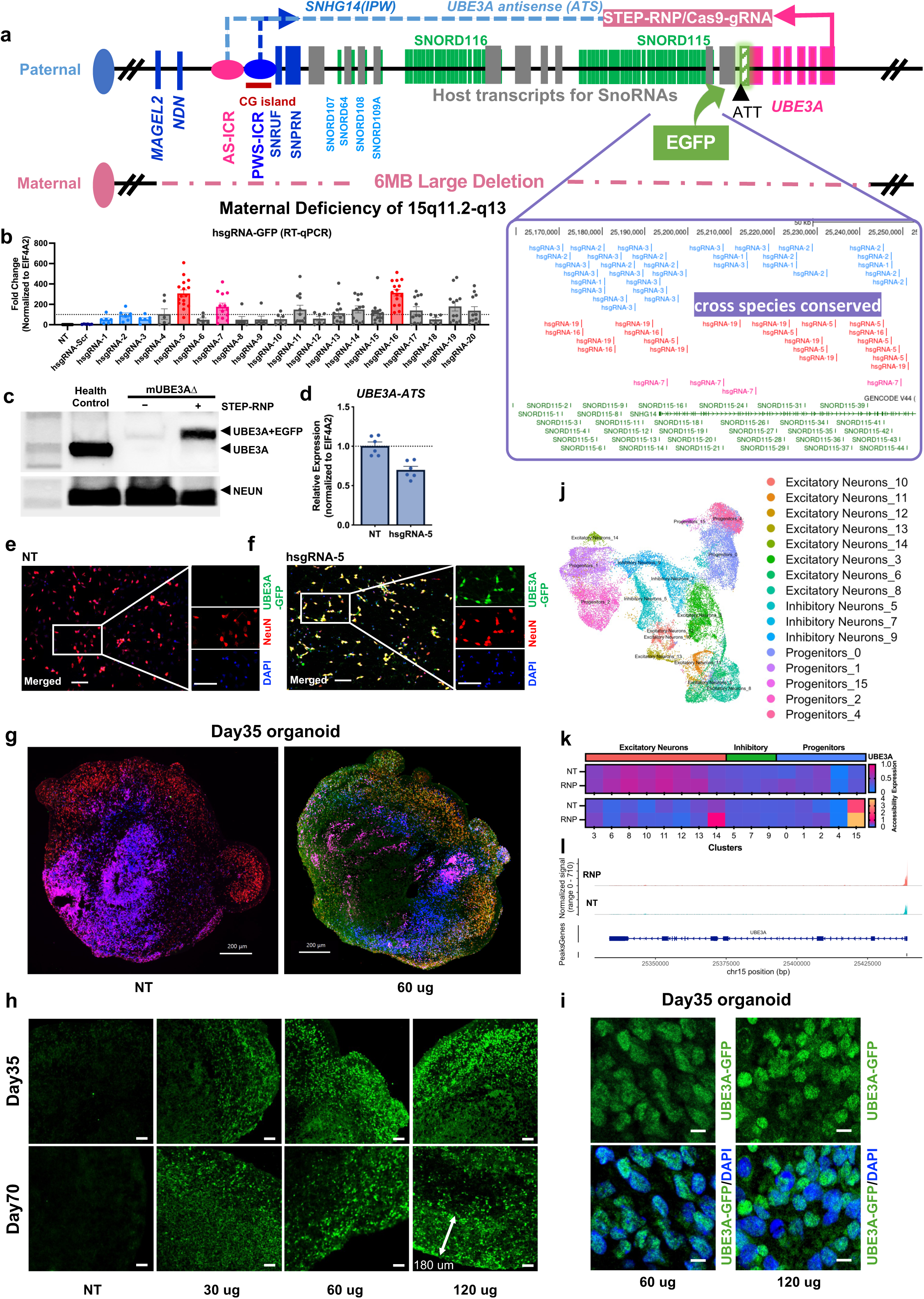
STEP-RNP editing restores UBE3A expression in induced neurons (iNs) and cortical brain organoids derived from Angelman syndrome (AS) iPSCs. **a.** Schematic of the paternal *UBE3A-EGFP* fusion iPSC line generated from an AS patient carrying a maternal *UBE3A* deficiency due to a 6 Mb 15q11.2–q13 deletion. EGFP was fused in-frame to the C-terminus of the paternal *UBE3A* coding sequence before the STOP codon. A series of human-specific sgRNA (hsgRNA) were designed to target a cross-species conserved region within the *UBE3A-ATS* locus. **b.** Quantification of *UBE3A-EGFP* expression by RT-qPCR screening 20 hsgRNA candidates in AS-iNs three days after incubation with STEP-RNP (15 µg/mL, 24 h incubation). hsgRNA-5, -7, and -16 exhibited the highest reactivation efficiency. **c.** Reactivation of paternal *UBE3A-EGFP* protein in AS-iNs following STEP-RNP-hsgRNA-5 treatment, confirmed by immunoblot. **d.** Reduction of paternal *UBE3A-ATS* transcripts in iNs treated with STEP-RNP-hsgRNA-5. **e-f.** Representative fluorescence images showing reactivation of *UBE3A-EGFP* in AS-iNs treated with STEP-RNP-hsgRNA-5 (f) compared to non-treated (NT) controls (e) three days after treatment (G5, 15 µg/mL, 24 h incubation; scale bar = 100 µm). **g.** Reactivation of paternal *UBE3A-EGFP* in cortical brain organoids derived from AS iPSCs treated with STEP-RNP-hsgRNA-5 (60 µg for 24 h incubation on day 28) and collected seven days later on day 35 (*n* = 6-8; scale bar = 200 µm). **h-i.** Dose-dependent reactivation of paternal *UBE3A-EGFP* in day 35 and day 70 cortical organoids treated with STEP-RNP-hsgRNA-5 at concentrations of 30, 60, and 120 µg (scale bar = 50 µm). **(h)** Top: day 35 organoids treated with STEP-RNP-hsgRNA-5 at day 28 for 24 hours. Bottom: day 70 organoids treated at day 63 for 24 hours, showing dose-dependent reactivation of *UBE3A-EGFP*. The reactivated *UBE3A-EGFP* signal extended up to ∼180 µm when 120 µg STEP-RNP was applied per 3 mL of culture medium (*n* = 6-8). **(i)** Enlarged view of day 35 organoids demonstrating higher *UBE3A-EGFP* expression in organoids treated with 120 µg compared to 60 µg STEP-RNP-hsgRNA-5 (scale bar = 10 µm). **j-k.** Single-cell Multiome (scRNA-seq + scATAC-seq) analysis of paired cortical organoids derived from AS iPSCs treated with STEP-RNP-hsgRNA-5 at day 28 and collected on day 35 and non-treated cortical organoids. STEP-RNP treatment significantly increased *UBE3A* expression across excitatory neurons, inhibitory neurons, and neural progenitor cell populations.

We used the sgRNA designer (Broad institute) to design 20 new hsgRNA candidates with high predicted efficiency using chr15:25163154-25260993(hg38) as a reference. Among them, 3 (hsgRNA1-3) were documented in a previously published study ^41^. Using AS-iPSC-iNs, we screened these 20 candidates in AS-iPSC-iNs and identified three (hsgRNA-5, hsgRNA-7, and hsgRNA-16) with the highest efficiency of *UBE3A-EGFP* reactivation as assessed by RT-qPCR (**Fig.5b and Extended Data 6c**). The editing efficiency was also confirmed by the reactivation of UBE3A and the reduction of UBE3A-ATS using Western blot and RT-qPCR analysis, respectively (**Fig. 5c,d**). Immunostaining performed three days post-treatment further confirmed the high efficiency of *UBE3A* reactivation for all three hsgRNAs paired with the G5 Cas9 variant (**Fig. 5e-f & Extended Data 6d).** None of the three candidates reduced the expression of surrounding genes *SNORD115*, *SNORD116*, *SNRPN* and *SNHG14/IPW* (**Extended Data 6e**). Overall, these data demonstrate that STEP-RNPs allow for efficient gene editing and *UBE3A* reactivation in human iPSC-derived neurons. We further characterized STEP-RNPs carrying hsgRNA#5 formulated with Cas9 variants of V3 Cas9, G5 Cas9, or G221 Cas9 that have minor differences in amino acid sequence. Among the three Cas9 variants, G5 Cas9 showed superior efficiency in which the paternal UBE3A-EGFP fusion protein was detectable in over 90% of the AS-iPSC-iNs 24 hours after treatment (**Extended Data 6f**). We therefore used the G5 Cas9 variants for the subsequent experiments.

To further evaluate the editing efficacy in human neurons, we performed STEP-RNP treatment of AS-iPSC-EGFP-derived cortical organoids using hsgRNA#5 and the G5 Cas9 variant. Highly efficient editing and reactivation of paternal UBE3A-EGFP were observed 7 days after a 24-hour incubation in both 35- and 70-day cortical organoids and the penetration of STEP-RNPs was dose dependent (**Fig. 5g-i**). Single nuclear RNA-seq and ATAC analyses of day 35 cortical organoids at 7 days after STEP-RNP treatment revealed significant reactivation of *UBE3A-EGFP* in both excitatory and inhibitory neurons as well as in neural progenitor cells (**Fig. 5j-k**), along with the increased chromatin accessibility of UBE3A gene by ATAC-seq analysis (**Fig. 5l**).

### On target editing profile and off-target assessment

We performed extensive evaluation of on-target and potential off-target events associated with STEP-RNP treatment using a combination of in silico, biochemical, cellular and molecular assays, in vitro in human iPSCs and iPSC-derived neurons and in vivo in the mouse brain (**Fig. 6a**). For the in vivo assessment, we performed in silico off-target predication with Cas-Offinder followed by amplicon sequencing and deep Illumina short read whole genome sequencing (300X WGS) followed by bulk RNA-Seq. For human iPSC-derived neurons, we followed recommendations of our FDA INTERACT meeting and performed in silico analysis and CHANGE-seq, followed by enrichment capture and deep short read, and PacBio long read sequencing to compare pre- (n=4) and post-STEP-RNP treated (n=4) iPSCs and iPSC-derived neurons.

**Fig 6.**
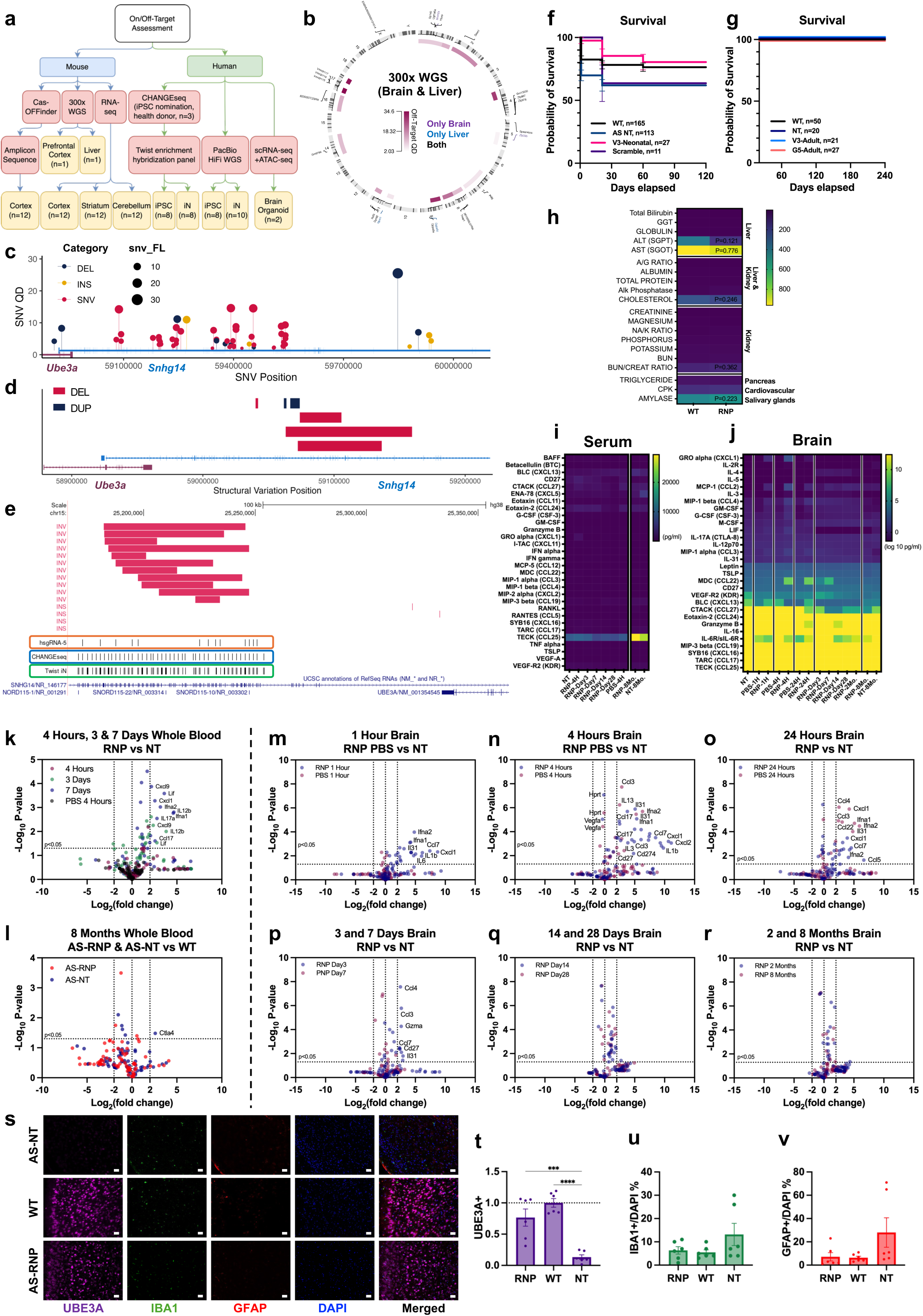
On- and off-target specificity, safety and immunogenicity of STEP-RNP delivery in iPSCs and mice. **a.** Workflow of on- and off-target assessment combining unbiased whole-genome sequencing (WGS), RNA-seq, and targeted enrichment using CHANGE-seq–nominated candidate sites. **b.** Extra deep (300×) WGS of cortex and liver tissues collected 30 days after IT STEP-RNP administration identified two off-target single-nucleotide variants (SNVs) in *Zfp33b* and *Mroh2a* exclusively in the brain, with no corresponding alterations in the liver. **c.** 300× WGS of prefrontal cortex 30 days after P1 IT STEP-RNP delivery revealed high on-target editing efficiency (indels: Deletion, *n* = 9; Insertion, *n* = 7; SNV, *n* = 46). **d.** Six structural variants (1.9 kb–64 kb) aligned with the targeted region were detected from the 300× WGS dataset. **e.** PacBio HiFi WGS in iPSC-derived neurons 3 days after a 24-h STEP-RNP incubation identified 11 on-target inversions (11-63 kb) and 4 insertions (21-322 bp). **f-g.** Survival analysis of WT and AS mouse cohorts following STEP-RNP treatment at neonatal (f) or adult (g) stages. P1-treated AS mice showed improved survival relative to untreated and scrambled controls, whereas all adult-treated groups survived through the study period. **h.** Serum chemistry profiles showed no significant differences in liver or kidney function parameters between STEP-RNP-treated and WT littermates six months after P1 IT administration. A mild, non-significant increase in AST levels was observed in both treated and untreated groups. **i-r.** Assessment of immune and inflammatory responses following STEP-RNP delivery. (i) *Serum protein profiling.* P21 WT mice were treated via IT with STEP-RNP-sgRNA-NC (scramble sgRNA), PBS, or untreated. A 64x multiplex protein-based immune panel measuring cytokines, chemokines, and growth factors revealed transient increases in several analytes (e.g., IL-22) at 4 hours after treatment with both STEP-RNP-sgRNA-NC and PBS compared with untreated (NT) controls. No significant changes were observed at 3, 7, or 14 days, although several factors were modestly elevated at day 28 (NT, *n* = 12; WT *n* = 6-8 each timepoint; PBS, *n* = 4). At 6 months post-P21 IT administration, no differences were detected between WT and STEP-RNP-AS-treated groups (*n* = 6). (j) *Brain protein profiling.* Analysis of the same immune panel in brain tissue showed transient upregulation of cytokines at 1, 4, and 24 hours post-RNP injection. The temporal pattern was nearly identical to that in PBS-injected animals, indicating a general injection-related response rather than a STEP-RNP-specific effect. All values returned to baseline by 14 days post-injection (*n* = 6). (k-l) *Whole-blood RNA profiling.* RNA-based 81x multiplex immune response panels applied to whole blood from P21 mice treated by IT with STEP-RNP-sgRNA-NC, PBS, or untreated showed no significant transcriptomic changes 3 days after injection. By day 7, 8 of 81 measured inflammatory or signaling genes were upregulated relative to NT controls, all of which returned to baseline by 8 months (NT, WT 4 h, WT 3 d *n* = 6; PBS *n* = 4). (m-r) *Brain RNA profiling.* Analysis of the same RNA-based 81x multiplex immune response panels in brain tissue revealed trends consistent with the protein assays. Both RNP- and PBS-injected mice displayed similar transient activation across cytokine, chemokine, and growth factor pathways. The RNP-treated group exhibited a slightly earlier onset of response (m), but by 4 h and 24 h post-injection, expression profiles were synchronized between RNP and PBS groups. By 14 days, all immune parameters had returned to baseline, indicating resolution of the transient inflammatory response (*n* = 6). **s-v.** Immunofluorescence staining (s) and quantification (t-v) of IBA1+ microglia and GFAP+ astrocytes AS mouse brain 8 months after STEP-RNP administration. STEP-RNP-treated AS mice showed reduced neuroinflammation compared with untreated controls (*n* = 6 per group, scale bar = 50 µm, * p<0.05, ** p<0.01).

On-target effects were identified via 300X WGS of mouse brain tissue. We identified 62 single nucleotide variants (SNVs) and 6 structural variations (SV) ranging from 1894 bp to 64 kb in the targeted region aligned to mouse chr7:58878550-60099925 (**Fig. 6c-d**). Similarly, high frequent (Allele Frequency = 0.421 ± 0.234) on-target events including SVs of microdeletion (DEL), microduplication (DUP) and inversion (INV) were identified in human iPSC-derived neurons using PacBio long reading sequencing (**Fig. 6e**). The detection of multiple structural variants is expected given that the hsgRNA is designed to target multiple sites within *UBE3A-ATS*. The number of on-target SVs detected may be underrepresented because of the average sequencing depth of the long read sequencing method. Hybrid capture using a custom-designed enrichment oligo panel followed by deep short-read sequencing (11,000X) of 4 pairs of pre- and post-STEP-RNP-treated iPSC-derived neurons (**Extended Data 7d-k**) revealed an on-target indel frequency of up to 53.04% within the chr15:25193313–25439024 region (**Extended Data 7l**). The presence of multiple hsgRNA target loci in the *UBE3A-ATS* probably contributed to the highly efficient editing but also created technical challenges in estimating editing efficiency per cell due to the complex nature of the editing outcomes. The indels and SVs likely disrupt and reduce the silencing function of *UBE3A-ATS,* which in turn results in the reactivation of *UBE3A* expression as described above.

To assess potential off-target events in mouse brains, we first examined the top 10 in silico predicted off-target sites for the mouse STEP-RNPs using the amplicon sequencing method described by AAV-mediated Cas9 genome editing ^41^. No off-target events were detected in the brain tissues of 6 mice treated with STEP-RNPs by IT administration at P1, compared to non-treated littermates. To perform the off-target assessment in an unbiased way, we performed short read WGS at 300X depth of brain and liver tissues from one of the same cohort of STEP-RNP treated mice. Because STEP-RNPs leak to non-brain tissues via IT administration, it was not feasible to design STEP-RNP treated and not treated tissues from the same mouse. Liver tissue was included because of high biodistribution of STEP-RNPs after IT delivery. Further, given the expected genome variability among WT mice, it was similarly not feasible to compare to 300X WGS from WT C57BL6/J mice due to a large number of mice that would be required due to the expected inter-variability of individual mouse genomes. Instead, we analyzed the 300X WGS from the STEP-RNP-treated mouse against a large number of reference mouse WGS data (∼30X) from the Mouse Genome Project (MGP) ^63,64^. We identified 20 variants; however, these were present in both brain and liver samples, suggestive of germline variants. Only two variants - in *Zfp33b* and *Mroh2a -* were unique to the brain. However, bulk RNA-seq of neocortex, striatum, and cerebellum tissues from 6 WT- and 6 STEP-RNP-treated mice via IT at P1 did not reveal significant transcriptomic changes, including of *Zfp33b* and *Mroh2a,* in the STEP-treated tissues (**Fig. 6b & Extended Data 7a-c**).

For the assessment of off-target events in iPSCs and iPSC-derived neurons, we performed Cas-Offinder ^65^, CHANGE-Seq followed by hybrid capture and deep next generation sequencing (NGS) for nominated sites ^66^, and PacBio long-reading sequencing. The CHANGE-Seq analysis in three iPSC lines from healthy donors nominated 4,422 potential sites, with 3.5% located in coding regions. These nominated sites were verified using a custom-designed enrichment oligo panel and capture (Twist), followed by 11,000 X deep NGS. We compared four STEP-RNP treated and non-treated iPSC-derived neuron lines and verified 269 shared off-target events from across the four lines at a saturated sequencing depth (**Extended Data 7d-k**). However, none of these sites reached an allele frequency (AF < 0.05). Altogether, our extensive analyses did not detect any significant off-target events associated with STEP-RNP treatment in mouse brains or in human iPSC-derived neurons.

### General safety and toxicity profile of STEP-RNPs

To assess the potential toxicity of our STEP-RNP formulation, we first evaluated the general health of treated mice, their weight and survival rates after STEP-RNP administration up to 120 (neonatal-treated) and 240 (adult-treated) days. STEP-treatment was not associated with decreased survival or weight in either the neonate or adult-treated group (**Fig.6f-g & Extended Data 5j**). The survival rate of AS-treated neonatal pups increased compared to NT and scramble sgRNA groups (p<0.05) and was comparable to WT. The observed reduced early survival rate across different groups is likely due to the increased infanticide associated with toe clipping and handling of newborns that is necessary for the experimental manipulations.

General toxicity was assessed using a serum-based mouse multiple organ function panel that covers hematological, hepatic, renal and other organ functions. No significant toxicity for organ functions was associated at 6 months post-STEP-RNP treatment of P1 AS mice via IT administration (**Fig.6h**). General histological analysis of the brain, liver, and other organs also did not reveal significant cellular changes associated with STEP-RNP treatment intrathecally using different histological methods including H&E (**Extended Data 8a-k**), Luxol Fast Blue (**Extended Data 8l-r**), and Masson’s Trichrome Staining (**Extended Data 8s-v**) of major organs after administration of STEP-RNPs. It is possible that a small amount of free STEP molecules may detach from the STEP-RNP complexes and bind endogenous proteins, such as serum proteins; however, STEP molecules are less likely to cause detrimental effects, as the compound shows no significant inhibitory activity on cytochrome P450 (CYP) enzymes or mutagenicity (data available upon request). The comparable behavioral profile (up to 6 months of age) between mice treated with STEP-RNPs-scramble sgRNA and non-treated mice further supports the safety profile of STEP-RNPs.

We next investigated both peripheral and central nervous system inflammation following IT administration of G5-formulated STEP-RNPs. Peripheral inflammatory and immune responses were assessed by profiling 64 cytokine, chemokine, and growth factor protein targets in mouse serum and brain tissues collected from 1 hour to 8 months post-IT administration. In serum, acute inflammation, observed at 4 hours post-STEP-RNP administration, was comparable to that following PBS administration and resolved to baseline levels after 3 days, although slightly increased levels of a few cytokines at day 28 were noted. By 6 months, no difference in the level of inflammatory markers was observed between WT and STEP-RNP-treated mice (**Fig.6i & Extended Data 9a**). In brain tissues, the results were generally comparable between non-treated and treated with PBS or STEP-RNPs at different time points (1h, 4h, 24h, 3d,7d, 14, 28, 60, 240d) (**Fig.6j and Extended Data 9b**). We also analyzed the expression of 80 RNA targets encoding cytokines, chemokines, and growth factors in whole blood and brain tissues from WT mice treated with STEP–RNPs intrathecally at P21. Consistent with the protein panel findings, only 8 out of 64 inflammatory markers showed increased expression in blood 7 days after injection, indicating limited inflammation (**Fig.6k)**. By 8 months post-IT administration, all cytokine, chemokine, and growth factor expression levels had returned to normal **(Fig.6l**). In brain tissues, the increased inflammatory markers in STEP-RNP treated group were observed 1h, 4h, and 24 h but significantly reduced by 3 and 7 days and normalized after 14 days (**Fig. 6m-r**). Immunofluorescent staining for IBA1 and GFAP at 1, 3, and 7 days and 8 months post-STEP-RNP administration revealed similar expression levels of IBA1^+^ microglia and GFAP^+^ astrocytes in treated and control mice, suggesting minimal neuroinflammation (**Fig. 6s-v & Extended Data 9c-g**). The number of IBA1^+^ microglia and GFAP^+^ astrocytes was higher in untreated AS mice than in wild-type and STEP-RNP-treated AS mice, consistent with their increase in expression due to spontaneous seizures ^67^. Interestingly, IBA1^+^ microglia and GFAP^+^ astrocytes are comparable between STEP-RNP-treated and wild-type mice (**Fig. 6u,v**). This result may indirectly support the reduced seizure susceptibility of STEP-RNP treatment in AS mice.

## Conclusion

One of the major challenges in translating CRISPR genome editing into clinical applications for neurogenetic disorders is the lack of delivery platforms that allow for efficient and safe brain-wide delivery. Current viral, peptide, and nanoparticle-based delivery methods have well-documented limitations, particularly for application to the brain in humans ^8,12–19,22–27^. Delivery of genome editing in RNP form offers a major advantage from a therapeutic perspective as it delivers active genome editing machinery directly and transiently, reducing safety concerns of Cas9 induced off target events associated with persistent transgene expression. However, the inability of RNPs to self-penetrate cells in the brain limits their clinical application ^8,12,13^.

Our approach to chemically engineering RNPs into STEP-RNP complexes overcomes this challenge. STEP-RNPs are only ∼13 nm in size; significantly smaller than the AAV vectors (∼25 nm) and nanoparticles (40-300 nm). This compact size, along with negative surface charge, allows for brain-wide penetration through IT and ICV administration at different developmental stages in mice. We have also observed brain wide distribution and editing via IV route delivery, although with lower efficiency. Given the appeal of IV administration in a clinical setting, additional characterization of this delivery route is warranted. We also show that STEP-RNPs efficiently edit human iPSC-derived 2D neurons and 3D cortical organoids. The high frequency of indels and structural variants induced by STEP-RNPs is the likely mechanism that inactivates the *UBE3A-ATS*, leading to the long-term reactivation of *UBE3A* from the paternal chromosome. Notably, UBE3A restoration in the mouse brain at various time points resulted in partial and/or complete rescue of neurobehavioral impairments in the AS *Ube3a* maternal deficiency mouse model. It is important to note that reactivating even 5-15% of UBE3A expression is expected to have a significant clinical impact in AS patients, based on studies of rare AS patients with mosaic imprinting mutations^68,69^.

Our findings demonstrate that STEP-RNPs overcome major delivery hurdles associated with traditional genome-editing platforms and present an innovative approach for delivery of genome editing to the brain. The cell-type specificity and cellular mechanism associated with STEP-RNPs remain to be further elucidated; however, our evidence demonstrates that STEP-RNPs are highly efficient in human and rodent Ai9 fibroblast, neurons, and NPCs at a minimum. We also observed editing in GFAP^+^ astrocytes via local CED administration; however, significant editing was not observed in GFAP^+^ astrocytes in Ai9 reporter mice. Due to the biallelic expression of UBE3A in astrocytes, we were unable to assess editing using UBE3A-YFP reporter mice. Future optimization of the STEP-RNP formulation in appropriate reporter systems is warranted to enhance its efficacy in non-neuronal populations. Additionally, biodistribution studies in larger animal models, such as non-human primates, as well as studies using GMP-grade STEP-RNPs are essential before moving into human applications.

Transcriptional disruption or inactivation of *UBE3A-ATS* is critical for un-silencing the paternal *UBE3A* gene ^36,37^; yet the underlying molecular mechanism remains incompletely understood. Our on-target analysis revealed frequent indels and structural variants including small deletions, duplications, and inversions across the *UBE3A-ATS* genomic region. It is technically challenging to precisely quantify the per–cell frequency of on–target events because of the multiplex targeting achieved by a single sgRNA and the complex mixture of editing outcomes across the UBE3A–ATS region. Nevertheless, our integrated approach combining short– and long–read DNA sequencing, bulk and single–cell RNA profiling, and in situ immunostaining of a UBE3A reporter-supports high editing efficiency in neurons following STEP–RNP delivery in human brain organoids and in mice, despite the fact that these methods may under-report the number of on-target events. It is important to note that in one prior AAV–Cas9 study, viral integration into the UBE3A–ATS region was detected at low frequency (∼2.8% of cells); however, this rate is unlikely to explain the strong reactivation observed^41^. Another study using a different sgRNA reported ∼2.1% AAV integration and ∼19.2% indels; the indel frequency correlated with the level of reactivation, but the underlying mechanism remains unknown^40^. We speculate that the frequent (∼40%) occurrence of diverse genomic alterations particularly small deletions (1.8 to 60 kb), duplications, and inversions induced by STEP–RNP editing could disrupt the continuity or structural integrity of *Ube3a–ATS*, thereby enabling reactivation of *Ube3a* expression. As in the above and other studies using AAV–Cas9 to target UBE3A–ATS, our work does not aim to uncover the precise mechanism by which various genomic alterations lead to reduced UBE3A–ATS and sustained paternal UBE3A reactivation. Hence, further investigation is warranted. Similarly, whether double–strand breaks (DSBs) and the kinetics of the DSB repair process induced by STEP-RNP editing in neurons may contribute to sustained editing effects should also be considered^70^.

A comprehensive safety evaluation of STEP-RNPs is beyond the scope of this proof-of-concept study for platform development. However, initial safety assessments suggest a promising profile, with no detrimental behavioral impact, no detectable off-target effects, low immunogenicity, and 100% survival at 8 months post-treatment. Our off-target assessments, while extensive, will be further refined in future studies with a focus only on human neurons as recommended by FDA INTERACT meeting. The excellent safety profile of STEP-RNPs is likely due to the unique chemical composition of the selected STEP chemical and the transient nature of Cas9/RNP delivery, as we show that most Cas9/RNPs are degraded and eliminated from cells within 24 hours. We anticipate that our novel STEP-RNP platform will significantly advance clinical management of AS because other potential treatment modalities such as ASO, Cas13, ATF, and small molecule based-therapies only provide the transient reactivation of *UBE3A*^19,38–41^. More importantly, given the adaptable and modular design of the STEP-RNP platform, it holds the potential to enable precise gene editing solutions across a wide range of neurogenetic diseases and other diseases.

## Supporting information

Extend Fig

## Acknowledgments

We thank the Couragene team, including Conrad Leung, Ying Xiao, and Min-Han Lin for technical support. We thank Daryl Klein for kindly support in protein binding analysis. We thank In-Hyun Park and Jonghun Kim for technical support on brain organoid culture. We thank Daniel DiMaio and Yuka Takeo for technical support on endocytosis investigation. We thank Allyson Berent, Jennifer Panagoulias, Pj Brooks, and Timothy Lavaute critical reading the manuscript and discussion

## Funding

This work was supported by the following grants:

National Institutes of Health grant UG3TR004713 (YHJ, JZ)

National Institutes of Health grant UG3/UH3NS115597 (JZ)

Foundation for Angelman Syndrome Therapeutics (FAST) (YHJ, XL).

## Author contributions

Conceptualization: XL, YX, YB, CEH, YHJ, JZ

Methodology: XL, YX, YB, JY, ZW, CQ, SC, ST, BDK

Investigation: XL, YX, YB, JY, ZW, JF, WCQ, GL, ZT, JT, GM, YL, VK, RC, BC, ZM,YM

Funding acquisition: YHJ, JZ, CEH

Project administration: YHJ, JZ, CEH

Supervision: YHJ, JZ

Writing - original draft: XL, YX, CEH, YHJ, JZ

Writing - review & editing: XL, CEH, YHJ, JZ

## Competing interests

YHJ and JZ are co-founders of Couragene, Inc and are affiliated with CourageAS Bio.. XL is a scientific consultant for Couragene.

## Data and materials availability

Bulk RNAseq, scRNA-seq & scATAC-seq, WGS, Twist Enrichment Panel Sequencing data, PacBio HiFi reads are available in Sequence Read Archive: SUB14607142. All additional data are available upon request.

## Materials and Methods

### STEP stability assay

The freshly synthesized STEP compound was stored in a -80°C freezer for two months before its purity was analyzed using HPLC. The analysis was conducted on a Shimadzu Nexera X3 LCMS-8050 or an equivalent instrument, utilizing an ACE Excel 2 C18-PFP column (2 μm, 3 × 150 mm). The mobile phase consisted of 0.1% TFA in water (Phase A) and 0.1% TFA with 5% THF in acetonitrile (Phase B). The HPLC method had a total runtime of 17 minutes, with the percentage of mobile phase B increasing from 25% to 99% before returning to 25%. The flow rate was set at 1.0 mL/min. All samples were dissolved or diluted in acetonitrile, and purity assessment was performed by analyzing the HPLC chromatogram peaks at UV 254 nm.

### Preparation of STEP RNP

1. **Preparation of Cas9/sgRNA RNPs:** In a typical synthesis, Cas9 and sgRNA were dissolved at 2 µg/uL and 1.2 µg/uL, respectively, in pH 7.4 PBS buffer. Cas9 solution (50 µL) and sgRNA solution (50 µL) were then mixed to form RNP (100 µL, with final concentrations of Cas9 at 1 µg/uL and sgRNA at 0.6 µg/uL).
2. **Preparation of STEP Cas9/sgRNA RNPs:** STEP molecules were dissolved at a concentration of 10 mM. In a typical synthesis, 2.5 µL of the STEP solution (10 mM) was added to the RNP solution (100 µL, containing Cas9 at 1 µg/uL) to achieve a STEP: Cas9 molar ratio of 20. The mixture was then incubated at room temperature for 30 minutes.
3. **Purification and Sterilization:** The RNP solution was washed with PBS buffer using an Amicon Ultra filter (Millipore, 10 kDa cutoff), concentrated to the desired concentration and sterilized by filtering through a 0.22 µm membrane for in vitro or in vivo use.

### Cas9 nucleases used for the studies

V3 Cas9 was purchased from Integrated DNA Technologies (IDT, Alt-R™ S.p. Cas9 Nuclease V3). G5 and G221 Cas9 are recombinant nucleases produced by Couragene Inc.

### SgRNA design and synthesis

sgRNAs were designed using the in-silico program provided by Broad Institute (https://portals.broadinstitute.org/gppx/crispick/public). sgRNAs used for the studies were purchased from Synthego.

### Ai9 Reporter Fibroblasts Culture

Ai9 reporter embryonic fibroblasts isolated from the Ai9 mouse (B6.Cg-*Gt(ROSA)26So ^tm^*^9^(CAG–tdTomato)*^Hze^*/J, Strain #:007909) were kindly provided by Prof. Jason Heaney at Baylor College of Medicine. The cells were cultured in high-glucose DMEM supplemented with 10% FBS, penicillin (100 U/mL), and streptomycin (100 μg/mL) and maintained at 37 °C with 5% CO2 in a humidified chamber.

### Analysis of Genome Editing Efficiency In Vitro Using Ai9 Reporter Fibroblasts Cells

Ai9 reporter fibroblasts cells were seeded in 48-well plates prior to treatment (5,000 cells in 0.5 mL medium per well). STEP Cas9/sgRNA-AB RNPs were added to each well at the desired concentration and incubated for 24 hours (sgRNA-A sequence: AAAGAAUUGAUUUGAUACCG; sgRNA-B sequence: GUAUGCUAUACGAAGUUAUU).

After the treatment, cells were stained with Hoechst 33342 (1 µg/mL) for 10 minutes and imaged using a fluorescence microscope (Keyence, BZ-X710). The genome editing efficiency was calculated as the percentage of tdTomato-positive cells relative to Hoechst-positive cells per imaging field ± SD (n = 3).

### Analysis of Genome Editing Efficiency In Vitro Using Ai9 Reporter NPCs

Neural progenitor cells (NPC) isolated from the embryonic Ai9 mouse were cultured in NPC medium: DMEM/F12 with GlutaMAX supplement, sodium pyruvate, 10 mM HEPES, penicillin and streptomycin, nonessential amino acid, 2-mercaptoethanol, B-27, N2 supplement, and growth factors bFGF and EGF (both 20 ng/ml as final concentration)^1^ . Ai9 reporter cells were seeded in 48-well plates prior to treatment (5,000 cells in 0.5 mL medium per well). STEP Cas9/sgRNA-AB RNPs were added to each well at the desired concentration and incubated for 24 hours. After the treatment, cells were stained with Hoechst 33342 (1 μg/mL) for 10 minutes and imaged using a fluorescence microscope (Keyence, BZ-X710). The genome editing efficiency was calculated as the percentage of tdTomato-positive cells relative to Hoechst-positive cells per imaging field ± SD (n = 10).

### Cell viability assay

Cells were seeded in 96-well plates prior to treatment (2,000 cells per well). STEP Cas9/sgRNA-AB RNPs were added to each well at the desired concentration and incubated for 48 hours. Cytotoxicity was tested by CellTiter-Glo® Luminescent Cell Viability Assay according to the protocol. Luminescence (RLU) was measured using Perkin Elmer Victor X2 Multilabel Microplate Reader. The relative cell viability (%) was expressed as [RLU]sample/[RLU]untreated × 100.

### Size and Zeta-Potential Measurement by Dynamic Light Scattering (DLS) Analysis

The hydrodynamic sizes of RNP and STEP-RNP were measured by dynamic light scattering (DLS) using a Malvern Nano-ZS (Malvern Instruments). RNP and STEP-RNP were diluted to 40 µg/mL with pH 7.4 PBS buffer before the measurement. The same solution was then loaded into a disposable capillary cell to measure the zeta-potential using the Malvern Nano-ZS.

### Transmission Electron Microscopy

RNP and STEP-RNP samples were diluted with PBS and applied to holey carbon-coated copper grids (Electron Microscopy Sciences). The grids were stained with one drop of NANO-W (Nanoprobes) for 1 minute and dried by air flow. Images were captured using a transmission electron microscope (FEI Tecnai TF20 TEM).

### Cellular Uptake Mechanism Study

Ai9 reporter cells were seeded in 48-well plates prior to treatment (5,000 cells in 0.5 mL medium per well). Cells were pretreated with various endocytosis inhibitors-PitStop (50 µM), Nystatin (100 µM), Amiloride (0.5 mM), or incubated at 4 °C for 30 minutes, then treated with STEP GFP-Cas9/sgRNA-NC RNP for 4 hours (Cas9 10 μg/mL). After treatment, cells were digested using 0.05% trypsin and harvested for flow cytometric analysis of GFP fluorescence (BD LSRII).

### Protein Binding Assay

Plasma (CD-1 mouse, BiolVT) were thawed and centrifuged at 3220 ×g for 5 minutes, and adjusted to pH 7.4 ± 0.1. Working solutions (400 μM) of STEP and control compounds were prepared by diluting stock solutions in DMSO. Loading plasma solutions (2 μM) were prepared by further diluting the working solutions in a blank plasma and mixing thoroughly. For ultracentrifugation, T₀ samples were prepared by mixing the loading plasma with PBS (1:1), adding stop solution, and storing at 2–8°C. F samples were prepared by pre-incubating the STEP containing plasma at 4°C for 30 minutes, followed by ultracentrifugation at 4°C, 155000 ×g for 4 hours. T₄.₅ samples were obtained by continuing the incubation of the pre-incubated plasma at 4°C for 4 hours. Samples were transferred to 96-well plates, mixed with PBS/matrix (1:1), and stop solution was added. The mixture was vortexed, centrifuged at 4000 rpm for 20 minutes, and 100 μL of supernatant was collected for LC-MS/MS analysis. Blank samples were prepared by mixing blank plasma with PBS and processed similarly to the other samples. For data analysis, %Unbound was calculated as (100 × F) / Mean of T₄.₅, %Bound as 100 - %Unbound₀. Here, F represents the analyte/internal standard ratio in the protein-free sample, T₄.₅ is the analyte/internal standard ratio after 4.5 hours, and T₀ is the analyte/internal standard ratio at time zero.

### Mini-Ames Assay for Mutagenesis Evaluation

The Mini-Ames assay was performed to evaluate the mutagenic potential of the test article STEP in the presence and absence of S9 metabolic activation mix. The assay utilized five bacterial tester strains: *Salmonella typhimurium* (TA98, TA100, TA1535, and TA97a) and *Escherichia coli* WP2 *uvrA* (pKM101), with histidine or tryptophan auxotrophs, respectively (Molecular Toxicology, Boone, NC). Fresh bacterial cultures and freshly prepared test article formulations were used. STEP was tested at eight dose levels (1.5–800 µg per well) in three replicate wells per dose, while the negative control (DMSO) was tested in six wells. The study also included strain-specific positive controls: 2-Nitrofluorene (TA98), Sodium Azide (TA100, TA1535), ICR-191 (TA97a), and MNNG (WP2 *uvrA*) for -S9 conditions, and 2-Aminoanthracene for all strains under +S9 conditions. The test system was plated using top agar mixed with the bacterial culture, the test or control articles, and either the S9 metabolic activation mix or phosphate buffer (-S9). The mixture was then overlaid onto minimal glucose agar plates and incubated at 37°C for ∼67 hours. Revertant colonies were automatically counted using the Sorcerer Colony Counter, except for TA1535, which was counted manually due to the low spontaneous rates. Cytotoxicity was assessed based on bacterial lawn thinning or a dose-dependent reduction in revertant counts, and precipitation was evaluated visually and microscopically. The test was considered positive if a ≥2-fold increase (TA97a, TA98, TA100, WP2 *uvrA*) or ≥3-fold increase (TA1535) in revertant colonies was observed compared to the solvent control, along with a dose-response trend.

### CYP Inhibition Evaluation in Human Liver Microsomes

The assay was conducted in vitro using human liver microsomes to evaluate the potential of STEP to inhibit cytochrome P450 (CYP) enzymes, which play a critical role in drug metabolism. Human liver microsomes (Cat. No. 452117, Corning) were pulled from the freezer to thaw on ice. Substrate working solution was transferred to corresponding wells, and PB buffer was transferred to Blank wells in incubation plate (20 µL / well). After that, 158 µL of microsomes working solution was transferred to all wells in incubation plate. STEP or positive control working solution (2 µL) was transferred to the wells, and vehicle was transferred to the wells for no inhibitor control. The final STEP concentrations were 0.05, 0.15, 0.5, 1.5, 5, 15, 50 µM. The positive controls were α-Naphthoflavone (3 µM), Sulfaphenazole (3 µM), N-3-benzyl-nirvanol (1 µM), Quinidine (3 µM), Ketoconazole (3 µM). Incubation plate was pre-warmed at 37.0°C for 10 min. After a 10 min, 20 µL of NADPH working solution was added to initiate the reaction, the incubation plate was incubated at 37.0°C for 10 min. The reactions were terminated by adding 400 µL of stop solution containing internal standards. The mixture was shaken for 10 min and centrifuged at 3220 ×g for 20 min. An aliquot of 200 µL of supernatant was removed and mixed with 100 µL of Ultrapure water. The samples were shaken for 10 min and injected for LC-MS/MS analysis for metabolite of substrate. Finally, XL fit was used to plot the percent of vehicle control versus the test compound concentrations, and for non-linear regression analysis of the data. IC50 values were determined using 3- or 4-parameter logistic equation. IC50 values were reported as “>50.0 µM” when % inhibition at the highest testing concentration (50.0 µM) is less than 50%.

### Intracellular Trafficking Study Using Hela Cells and Ai9 NPCs

HeLa S3 cells were provided by Dr. Daniel DiMaio’s laboratory at Yale University School of Medicine and cultured at 37°C with 5% CO2 in Dulbecco’s modified Eagle’s medium (DMEM) supplemented with 20 mM HEPES, 10% fetal bovine serum (FBS), L-glutamine, and 100 units/mL penicillin-streptomycin (DMEM10). Hela cells were thawed 12-16 hours prior to STEP-RNP administration. Ai9 NPCs were seeded to coated plates 2-3 days prior to STEP-RNP administration. STEP-RNP-scramble sgRNA (GCGAGGUAUUCGGCUCCGCG) at a concentration of 10 μg/mL Cas9 was pre-mixed with culture medium and added to the cells at specific time points for the designated period. Following treatment, the medium was removed, and the cells were washed with PBS, then fixed with 4% formaldehyde solution in PBS for 15 minutes. Subsequently, cells were permeabilized and blocked simultaneously using 0.1% saponin in DMEM-10 (10% FBS culture medium) for 30 minutes. Primary antibody incubation was performed overnight at 4°C using 0.1% saponin in DMEM-10 for dilution. Secondary antibody incubation was carried out at room temperature for 1 hour.

### UBE3A-EGFP Fusion Reporter IPSC Line Generation and Culture

hIPSCs from Angelman Syndrome (AS) patient with a 6.21 Mb maternal deletion on 15q11.2-13.1 were generated at the Yale Stem Cell Core facility using CytoTune-iPS 2.0 sendai Reprogramming kit (Thermo-Fisher, Catalog # A16517) following manufacture protocol. The Derived iPSC clones were confirmed for clarence of residue virus, pluripotency by forming teratoma with three gem layers and genome stability using aCGH analysis. Clone number 3 was used for generating paternal UBE3A-GFP reporter line. The hIPSCs were cultured on Growth Factor Reduced (GFR) Matrigel (Corning, 354230)-coated 6-well plates and fed daily with mTeSR™ Plus medium (Stemcell Technologies, 05825). Cells were dissociated into single cells using ACCUTASE™ (Stemcell Technologies, 07920) for electroporation when they reached approximately 70-80% confluency.

For transfection, Cas9 protein (IDT, Alt-R^®^ S.p. Cas9 Nuclease V3) and sgRNA (IDT, Alt-R™ CRISPR-Cas9 sgRNA) were mixed in vitro to form an RNP complex, which was co-electroporated with EGFP plasmid donor flanked with 5’- and 3’-homologous arms to stop codon of *UBE3A* gene into approximately 20,000 hIPSC single cells using the Amaxa 4D-Nucleofector^®^ X Unit (Lonza, AAF-1003X). The electroporated hIPSCs were plated at low density onto GFR Matrigel-coated 6-well plates and cultured for about 10 days to form single-cell colonies. The colonies were picked as duplicates: half of the clone was placed in a well of a GFR Matrigel-coated 48-well plate for continued culture, and the other half was used for positive clone screening by PCR. The positive clones identified through screening were subsequently subcloned to ensure purity, and Sanger sequencing confirmed them as correctly edited clones. Thereafter, these AS patient-derived hIPSCs, with in-frame EGFP fusion to the C-terminus of the paternal *UBE3A* gene before the STOP codon (AS-hIPSCs-EGFP), were maintained in mTeSR™ Plus medium.

### Lentiviral Package

Lenti-X™ 293T cells (Takara, 632180) were transfected with the plasmid vectors of pLV-TetO-hNGN2-Puro (Addgene, #79049) and Fudelta GW-rtTA (Addgene, #19780) using Lenti-X Packaging Single Shots (VSV-G) (Takara, 631275) according to the manufacturer’s instructions. The generated lentivirus was purified using the Lenti-X™ Maxi Purification Kit (Takara, 631233) and the buffer was switched to mTeSR™ Plus medium using disposable PD-10 Desalting Columns (Sigma, GE17-0851-01). Lentiviral titers were then determined using the Lenti-X™ qRT-PCR Titration Kit (Takara, 631235) and Lenti-X™ GoStix™ Plus (Takara, 631280).

### Transduction and Generation of NGN2 Inducible hIPSCs

AS-hIPSCs-EGFP were cultured in GFR Matrigel-coated 6-well plates and fed with mTeSR™ Plus medium until they reached approximately 70-80% confluency. On day 1, a master mix medium containing pLV-TetO-hNGN2-Puro lentivirus and Fudelta GW-rtTA lentivirus (both titers >10^9^) at 1 µl/ml medium was prepared and used to feed the cells for 48 hours. On day 3, the cells were either passaged for expansion (for less than 10 passages) or used for neuronal differentiation.

### AS-hIPSCs-EGFP-iNeurons Differentiation and Culture

The standard NGN2 overexpression iNeuron protocol was modified to facilitate STEP-RNP application ^2^. Briefly, AS-hIPSCs-EGFP cells were maintained in mTeSR™ Plus medium until they reached approximately 70-80% confluency. On day 1, the cells were fed with KSR medium containing doxycycline (dox; 2 µg/ml) to induce NGN2 expression, along with LDN-193189 (100 nM), SB431542 (10 µM), and XAV939 (2 µM) to enhance neuronal maturity. On day 2, the cells were fed with a 1:1 ratio of KSR medium, puromycin (puro; 5 µg/ml), and dox (2 µg/ml) to select for transduced cells. On day 3, the medium was changed to N2B medium supplemented with B27 (1:100), puro (5 µg/ml), and dox (2 µg/ml). On day 4, the cells were fed with full NBM medium containing NBM, B27 (1:50), BDNF/GDNF/CNTF (1:1000), puro (5 µg/ml), and dox (2 µg/ml), with or without STEP-RNP. On day 5, a complete medium change with full NBM medium was performed if the STEP-RNP was applied on day4. The cells were collected on day 7 for subsequent experiments.

### AS-iNeurons Immunostaining and Imaging

AS-iNeurons were maintained in NBM medium until STEP-RNP administration. STEP-RNP with different sgRNAs was pre-mixed with culture medium at a concentration of 10 μg/mL Cas9. Following treatment, the medium was removed, and the cells were washed with PBS, then fixed with 4% formaldehyde solution in PBS for 15 minutes. Subsequently, cells were permeabilized with 0.2% Triton X-100 in PBS for 15 minutes, followed by blocking with blocking buffer (CST, 12411) for one hour. The cells were then incubated with primary antibodies overnight at 4°C, followed by secondary antibody incubation for one hour at room temperature. Finally, the cells were mounted with DAPI Fluoromount-G® (SouthernBiotech, 0100-20).

### Cortical Brain Organoid Culture

Following established protocol^3^, AS-iPSCs were dissociated into a single-cell suspension and seeded into an ultra-low attachment 96-well plate at a density of 9,000 cells per well in 150 µL of induction medium supplemented with SB431542, LDN193189, and XAV939 on day 0 to initiate embryoid body (EB) formation. Induction medium was refreshed daily until day 10. On day 10, EBs were transferred to ultra-low-attachment 6-well plates for spinning culture at 80 rpm in neural differentiation medium supplemented with brain-derived neurotrophic factor (BDNF) and ascorbic acid, without vitamin A. Medium changes were performed every other day until day 18. From day 18 onward, organoids were cultured in neural differentiation medium containing BDNF and ascorbic acid to promote neural maturation, with medium refreshed every four days.

### Mice

Wild type C57BL/6J mice (Strain #:000664) and Ai9 reporter mice (B6.Cg-*Gt(ROSA)26So ^tm^*^9^(CAG–tdTomato)*^Hze^*/J,Strain #:007909) were obtained from the Jackson Laboratory^4^. *Ube3a^m+/p-YF^*^P^ (B6.129S7-*Ube3a^tm2Alb^*/J, Strain #:017765) ^5^ and *Ube3a^m-/p+^* (B6.129S7-*Ube3a^tm1Alb^*/J, Strain #:016590) ^6^ mice were generated at Baylor College of Medicine and maintained in Dr. Jiang’s lab at Yale University School of Medicine. *Ube3a^m-/p+^* and *Ube3a^m+/p-YF^*^P^ mice have been maintained on pure C57BL/6J background for more than 20 years. *Ube3a^m-/p+^* mice were produced from breeding between WT C57BL/6J male and *Ube3a^m+/p-^* female mice. Mice were housed of 4-5 per cage in pathogen-free mouse facility with free access to food and water, on a 12-hour light/dark cycle, at the ambient temperature of 20-22°C and humidity of 30-70%. An equal number of male and female mice were used for all experiments. All procedures were performed following the approved animal protocol by the Yale University School of Medicine Animal Care and Use Committee. Experimenters were blinded to the genotypes and treatments as experimental conditions permitted.

### In Vivo Genome Editing Analysis in Ai9 Brain via Convection-Enhanced Delivery

Ai9 mice (8-12 weeks of age, male/female) were anesthetized with a ketamine/xylazine mixture. After disinfecting the mouse head with povidone iodine and ethanol, a midline scalp incision was made, and a 1 mm hole was drilled on the right side of the skull, 2.5 mm lateral and 1 mm anterior to bregma. A Hamilton syringe with silica tubing was inserted 4.5 mm deep into the mouse striatum, followed by a 1 mm withdrawal. STEP Cas9/sgRNA-AB RNP (Cas9 10 μg/mL) was infused at a rate of 1.0 μL/min for 7 minutes. Following the infusion, the syringe remained in position for 10 minutes. The hole was sealed with bone wax, and the scalp was sutured. The mouse was injected with buprenorphine and back to home cage.

### Procedures of Intrathecal (IT), Intracerebroventricular (ICV), and Intravenous (IV) Injection of STEP-RNP

IT, ICV and IV administration of STEP-RNP were performed following published protocols ^7–9^. Briefly, the neonatal mice were cryo-anesthesia, adult were anesthesia with Isoflurane. Briefly, neonatal mice were anesthetized using cryo-anesthesia, while adult mice were anesthetized with isoflurane.

For IT injection, the needle was gently inserted and tilted slightly to a 70-80° angle at the intersection of indentation, keeping the syringe in a central sagittal plane. The angle was gradually reduced to approximately 30° as the needle touched the bone, then the needle was slipped (about 2 mm) into the intervertebral space. Up to 10 µL of STEP-RNP was slowly injected over 60 seconds for neonates, and up to 20 µL was injected into adults over 2-3 minutes. The needle was retained for 10-20 seconds post-injection to prevent leakage. The needle was withdrawn with gentle rotation to avoid leakage. In neonatal pups, the cerebellum turning green indicated correct needle placement, while in adults, a tail flip confirmed the correct location of the needle tip.

For ICV injection, the needle was inserted perpendicularly to the skull surface into the lateral ventricle, located approximately 0.8-1 mm lateral from the sagittal suture, halfway between lambda and bregma. A total of 2 µL of STEP-RNP was injected into each lateral ventricle over 2 minutes.

For IV injection in neonatal pups, the temporal vein, located just anterior to the ear bud, was used. A total of 10 µL of STEP-RNP was injected into the temporal vein over 2 minutes.

### Open Field Activity Test

To assess locomotor activity, mice will be tested on open field at 3-4 weeks of age, and a repeated test at 7-8 weeks. Mice will be individually placed in a 45 × 45 cm square open field and allowed to explore for 60 min. The total travel distance and movement per 5 minutes of each mouse in the open arena, and center zone, will be recorded by camera (Noldus Wageningen) connected to the EthoVision software (Noldus Wageningen).

### Rotarod

Motor performance on the rotarod will be assessed in acquisition session and retest session after training. Accelerating paradigm with 4 consecutive 5-min trials separated by an inter-trial interval of 60 minutes. In the accelerating paradigm, the rod began at 4 rpm and slowly increased to 40 rpm over the 5 minutes test. In the steady-state paradigm, the rod was maintained 16 rpm. After 48 hours break, two additional trials of both accelerating and steady-state paradigm will be conducted during the retest session. Latency to fall will be recorded, or a maximum of 300 second will be entered if the mouse remained on the rod for the duration of the test.

### Marble Burying

In the marble burying assay, each test was conducted in a 26 x 48 x 20 cm rat cage with a filter-top cover. Approximately 5 liters of fresh autoclaved bedding were evenly distributed to a depth of 5 cm. Twenty standard glass marbles, each 15 mm in diameter and weighing 5.2 g, and varying in style and color, were systematically placed on the flat surface of the bedding. Mice were allowed to explore the arena for 30 minutes. Post-exploration, a marble was scored as buried if at least half of its surface area was covered by the bedding.

### Novel Object Recognition Memory

Mice will be examined for short- (STM) and long-term memory (LTM). Training and testing will be conducted over 5 minutes in three phases: training, a test for STM at 30 minutes, and for LTM at 24 hours. At training, mice will be exposed to a pair of identical objects (2 x 2 x 3 cm in size) in a white Plexiglas arena (41 x 18 x 30 cm). These objects constituted the “familiar” objects for the tests. In the STM and LTM tests, one of the two familiar objects will be replaced with a novel object of similar dimensions to the training object but with different colors, patterns, and shapes. The durations of object contacts will be obtained.

### Fear Conditioning

Mice will be conditioned and tested for contextual and cued fear under ∼100 lux illumination in Med Associates chambers. On day 1 the mouse will be placed into the chamber for 2 minutes, after which a 72-dB 12-kHz tone (conditioned stimulus, CS) will be presented for 30 seconds, which co-terminated with a 2 seconds 0.4-mA scrambled foot-shock (unconditioned stimulus, UCS). This pairing will be repeated two additional times with an inter-stimulus interval of 60 seconds. Mice will be removed from the conditioning chamber to the home cage 1 minute after conditioning. For context testing on day 2, animals will be returned to the conditioning chamber for 5 minutes in the absence of the CS and UCS. For cued testing on day 3, the dimensions, texture, and shape of the conditioning chamber will be modified. Mice will be introduced into the chamber for 2 minutes, after which the CS will be presented for 3 minutes. For all tests, behavior will be videotaped and scored for freezing behavior by FreezeScan software (Cleversys, Reston VA).

### Seizure Induction

Seizure susceptibility was evaluated following a modified flurothyl-induced seizure paradigm ^10^. Briefly, a 10% solution of flurothyl [bis (2,2,2-trifluoroethyl) ether, Sigma] was diluted with 95% ethanol and infused onto a gauze pad at a rate of 10 mL/min via a syringe pump into an induction chamber. The mice were exposed for 5 minutes following a 1-minute period of free exploration. The interval from the initiation of flurothyl administration to the first occurrence of either 1) myoclonic seizures: characterized by abrupt, transient muscle contractions of the neck and body; or 2) tonic seizures: marked by a prolonged loss of posture control lasting over two seconds with concurrent trunk rigidity was recorded. The experiments were carried out in a well-ventilated cabinet to mitigate exposure to flurothyl vapors. The latency periods for myoclonic and tonic seizures in each group were compared.

### RNA Isolation and RNA Integrity Assessment

Mouse brain tissues were snap-frozen in liquid nitrogen immediately after dissection and stored at -80°C thereafter. Total RNA was isolated from 20 mg frozen tissues, using NucleoZOL™ (Takara Bio, 740404.200) and NucleoSpin® RNA set for NucleoZOL™ (Takara Bio, 740406.50) following the manufactures specifications, followed by rDNase Set (Takara Bio, 740963) to digest DNA, and NucleoSpin^®^ RNA Clean-up XS (Takara Bio, 740903) for RNA repurification. RNA purity (260/280, 260/230) and concentration were measured on NanoDrop™ 2000/2000c Spectrophotometers. RNA integrity number (RIN) was assessed using Agilent 2100 Bioanalyzer system. NucleoSpin^®^ TriPrep (Takara, 740966.50) and NucleoSpin® RNA/Protein Kit (Takara Bio, 740933.50) were used for DNA/RNA/Protein and RNA/Protein parallel isolation instead of the NucleoZOL™ for AS-hIPSCs-iNeurons, and when the mouse gDNA is needed for the WGS.

### Real-time Quantitative Polymerase Chain Reaction (RT-qPCR)

Two μg of total RNA was reverse transcribed into cDNA templates using RNA to cDNA EcoDry™ Premix kit including both random hexamer and oligo(dT)_18_ primers (Takara Bio, 639548). KAPA SYBR^®^ FAST qPCR Master Mix (2X) Universal (Kapa Biosystems, KK4602) was used for qPCR reactions with 18 ng of cDNA as template input. The following program on CFX96 Touch Real-Time PCR Detection System (BIO-RAD) was used: 3 minutes at 95°C for enzyme activation, followed by 40 cycles of denaturation (95°C, 3 seconds) and annealing, extension, data acquisition (60°C, 30 seconds), followed by dissociation and holding at 4°C.

### Western Blot

Whole cell lysates were extracted from mouse brain tissue using the NucleoSpin® RNA/Protein Kit (Takara Bio, 740933.50). Protein concentrations were quantified using the Protein Quantification Assay (Takara Bio, 740967.250). The samples were then mixed with 4x Laemmli buffer (Bio-Rad, 1610747) and heated at 98°C for 5 minutes to denature the proteins. Subsequently, proteins were loaded onto 4–20% Mini-PROTEAN^®^ TGX Stain-Free™ Protein Gels (Bio-Rad, 4568094) for electrophoresis. For immunodetection, the gels were incubated with primary antibodies overnight at 4°C, followed incubation with secondary antibodies an hour at room temperature.

### Mouse Tissue Collection and Immunostaining

Despite the need to split brain samples for molecular analysis (DNA/RNA/Protein) and immunostaining, all other samples, including whole-body samples, underwent perfusion with PBS followed by 4% paraformaldehyde (PFA, Santa Cruz Biotechnology, sc-281692). Samples were post-fixed overnight in 4% PFA at room temperature, then immersed in 70% ethanol before being submitted for paraffin embedding at Yale Pathology. The samples were sectioned at 5 µm thickness.

For immunofluorescence staining, the tissue sections were heated at 56°C for an hour, followed by deparaffinization with xylene (VWR, 89370-088) and rehydration. Antigen retrieval was conducted using citrate solution (Diagnostic Biosystems, 99990-096) for 20 minutes at 98°C, then cooled to room temperature. Tissue sections were permeabilized with 0.2% Triton X-100 in PBS for 15 minutes, followed by blocking with 5% donkey serum (Jackson ImmunoResearch, 017-000-121) for an hour. Sections were incubated with primary antibodies at 4°C overnight, followed by secondary antibody incubation for an hour at room temperature. Finally, the sections were mounted with DAPI Fluoromount-G^®^ (SouthernBiotech, 0100-20).

### Single Cell RNAseq

Cortical tissues (100 mg per sample) were collected from *mUbe3a*-YFP, *pUbe3a*-YFP P1 RNP–injected, and *pUbe3a*-YFP P21 RNP–injected mice at 52 days of age. Samples were snap-frozen in liquid nitrogen and stored at −80 °C until processing. Single-nucleus isolation and 10x Genomics single-nucleus RNA-seq library preparation were performed at the Yale Center for Genome Analysis (YCGA) using the GEM-X Universal 3′ Gene Expression v4 kit and Chromium GEM-X Single Cell 3′ Chip Kit v4, according to the manufacturer’s instructions. Raw sequencing data were processed with Cell Ranger (v9.0.1, 10x Genomics) using default parameters.

### Single Cell Multiome ATAC and Gene Expression

AS-brain organoids were treated with STEP-RNP on Day 28 for 24 hours and collected on Day 35, along with untreated Day 35 brain organoids. Samples were submitted to the Yale Center for Genome Analysis (YCGA) for single-nucleus isolation and 10x Multiome library preparation, following the manufacturer’s protocol. Raw sequencing data were processed using Cell Ranger ARC (v2.0, 10x Genomics) with default parameters.

### Gene Expression Analysis

Processed data were integrated and analyzed using Seurat (v4.0) in R (v4.4.0). Quality control steps included the removal of cells with fewer than 1,000 detected genes and genes expressed in fewer than five cells. Feature counts were normalized using total counts and scaled by a factor of 10,000. Highly variable genes (HVGs) were selected using variance-stabilizing transformation (VST), where genes were binned based on their mean expression, and the top 2,000 most variable genes were used for downstream analysis. Principal component analysis (PCA) was performed using the identified HVGs, and cell clustering was conducted using the first 30 principal components (PCs). Canonical correlation analysis (CCA) was applied to identify pairwise anchor correspondences across single-cell transcriptomes, enabling integration into a shared space. Gene expression values were scaled across all integrated cells before PCA. Using the first 30 PCs, cells were projected into a two-dimensional UMAP space, and clusters were identified accordingly. Differentially expressed genes (DEGs) in each cluster were determined using a fold-change threshold of 1.25 and a p-value < 0.05 (two-sided t-test).

### Cell Type Annotation

Cells were first classified into three major types based on marker gene expression: excitatory neurons, inhibitory neurons, and progenitors. Within each major group, further clustering was performed using gene ontology enrichment analysis, leading to the identification of 16 distinct subclusters. **(1) Excitatory neurons** included multiple subtypes, such as Layer 5-6 glutamatergic neurons (near-projecting and corticothalamic-projecting, CL0000679), Layer 2 glutamatergic neurons (LAMP5+ KCNG3+), Layer 5 intratelecephalon-projecting glutamatergic neurons, and those associated with angiogenesis (ADGRL4+). Additional subtypes included LINC00507+ GLRA3+ Layer 2, RORB+ LAMA4+ Layer 3-5 glutamatergic neurons, and neurons identified in the amygdala. **(2) Inhibitory neurons** were predominantly Sst+ GABAergic neurons (CL0000617), including a fetal brain-associated subtype, as well as Vip+ GABAergic neurons. **(3) Progenitors** included immune-related neural progenitors, cortical cingulate progenitors, and proliferating NK/T progenitors (CL0000542) in the prefrontal cortex, along with proliferating basal progenitors (CL0000646).

### Chromatin Accessibility Analysis

Chromatin accessibility data were processed using Cell Ranger ARC (v2.0, 10x Genomics), which performs read alignment, deduplication, and peak calling. The output fragment files were analyzed using Signac (v1.7.0, an extension of Seurat for scATAC-seq analysis). Peaks were filtered based on read depth and accessibility scores, and TF-IDF normalization was applied using RunTFIDF to correct for sequencing depth variability. Dimensionality reduction was performed using singular value decomposition (SVD) on the TF-IDF matrix, with the first 2–50 latent semantic indexing (LSI) components used for clustering and integration with gene expression data. To associate open chromatin regions with gene expression, co-accessibility scores were calculated using LinkPeaks, correlating ATAC-seq peaks with differentially expressed genes. Chromatin accessibility patterns were integrated with RNA-seq data to refine cell-type identities.

### Immune Profiling Panels

**For serum immune protein profiling,** 100 μL of serum was collected from each mouse and stored at −80 °C until analysis using the ProcartaPlex™ Mouse Immune Response Panel, 64-plex (ThermoFisher, EPX640-20064-901). Serum samples from each mouse were processed in triplicate.

**For brain immune protein profiling,** one hemisphere of each mouse brain was collected, snap-frozen in liquid nitrogen, and stored at −80 °C until processing. Tissues were homogenized in Cell Lysis Buffer (EPX-99999-000) supplemented with 1 mM PMSF Protease Inhibitor (ThermoFisher, 36978) and Halt™ Protease and Phosphatase Inhibitor Cocktail (ThermoFisher, 78441), using 500 μL buffer per 100 mg of tissue and a 5 mm stainless-steel bead (Qiagen, 69989) at 50 Hz for 8 min. Homogenates were centrifuged at 16,000 × g for 10 min at 4 °C, and supernatants were collected. Total protein concentrations were measured with the Pierce™ Dilution-Free Rapid Gold BCA Protein Assay Kit (ThermoFisher, A55861) according to the manufacturer’s instructions, adjusted to 10 mg/mL in 1× PBS, and analyzed using the ProcartaPlex™ Mouse Immune Response Panel, 64-plex (ThermoFisher, EPX640-20064-901). Each brain sample was processed in duplicate.

**For whole-blood RNA profiling, 1**00 μL of blood from each mouse was collected into 280 μL of PAXgene™ Blood RNA reagent (VWR, 77776-026) and stored at −80 °C. RNA was extracted using the QuantiGene™ Sample Processing Kit (ThermoFisher, QS0110), followed by analysis with the QuantiGene™ Plex Assay Kit (ThermoFisher, QP1013) and QuantiGene™ Plex Mouse Immune Response Panel, 80-plex (ThermoFisher, QGP-180-20064). Samples were processed in duplicate alongside standard RNA provided by ThermoFisher.

**For brain RNA profiling,** one hemisphere of each mouse brain was collected, snap-frozen in liquid nitrogen, and stored at −80 °C until RNA extraction. Total RNA was isolated using NucleoZOL™ (Takara, 740404.200) and the NucleoSpin® RNA Set for NucleoZOL™ (Takara, 740406.50), treated with DNase using the rDNase Set (Takara, 740963), and further purified with the NucleoSpin® RNA Clean-up XS kit (Takara, 740903.50) according to the manufacturer’s instructions. Purified RNA was adjusted to 300 ng in 20 μL and directly processed using the QuantiGene™ Plex Mouse Immune Response Panel, 80-plex (ThermoFisher, QGP-180-20064). RNA samples from each mouse were analyzed in duplicate alongside standard RNA provided by ThermoFisher.

The Luminex™ 200™ Instrument System (ThermoFisher, APX10031) was used for both protein and RNA assay detection, and the data were analyzed on the ThermoFisher Connect Platform.

### Amplicon Sequencing

Genomic DNA was extracted from mouse brain tissue using NucleoSpin^®^ Tissue kit (Takara, 740952.50), following the manufacturer’s protocol. KAPA SYBR^®^ FAST qPCR Master Mix (2X) Universal (Kapa Biosystems, KK4602) was used for PCR reactions with 18 ng of cDNA as template input. The following program on CFX96 Touch Real-Time PCR Detection System (BIO-RAD) was used: 3 minutes at 95°C for enzyme activation, followed by 40 cycles of denaturation (95°C, 3 seconds) and annealing, extension, data acquisition (60°C, 30 seconds), followed by dissociation and holding at 4°C. The target specific PCR primers are shown in supplementary Table, with following cap added: 5’-ACACTCTTTCCCTACACGACGCTCTTCCGATCT [target specific primer forward]-3’ and 5’-GTGACTGGAGTTCAGACGTGTGCTCTTCCGATCT [target specific primer reverse]-3’. The amplicons were purified using NucleoSpin® Gel and PCR Clean-Up (Takara, 740609.50), and quantified using Qubit 4 and Qubit dsDNA HS Assay Kit (Invitrogen™, Q32854). 200 ng of genomic DNA per mouse (pool amplicon of 10 off-target regions) were submitted to Yale Center for Genome Analysis (YCGA) for library preparation. Sequencing libraries were prepared using the Illumina Nextera XT DNA Library Preparation Kit, according to the manufacturer’s instructions. The libraries were then sequenced on the Illumina MiSeq platform using MiSeq Reagent Kit v3, generating 1 million 100bp paired-end reads.

The raw sequencing reads were processed to remove adapters and low-quality bases using Cutadapt^11^. High-quality reads were aligned to the reference genome (GRCm39) using BWA-MEM with default parameters^12^. Post-alignment, BAM files were sorted, and duplicates were marked using Picard tools^13^. Variants were called using GATK HaplotypeCaller ^14^, generating VCF files for each sample. The VCF files were filtered to focus on the off-target regions using bcftools (v 1.16)^15^.

### RNA-Seq Data Production and Processing

RNA-seq production was performed at Yale Center for Genome Analysis (YCGA) using the standardized protocol. Illumina bulk RNA-Seq raw data in FASTQ format after quality control and filtering with fastp^16^, were aligned to GRCm39 for mouse sequences using STAR 2.7.11b^17^. Gene expression counts and DEXSeq-counts were calculated using FeatureCount^18^ for further gene expression analysis.

### Whole Genome Sequencing (WGS) Analysis

Mouse gDNA was sequenced at Yale Center for Genome Analysis (YCGA) at 300X coverage using Illumina platform. Raw sequencing reads were generated in FASTQ format. The quality of raw sequencing reads was assessed using fastp ^16^. Adapter trimming and quality filtering were performed with Trim Galore to ensure high-quality reads. High-quality reads were aligned to the reference genome (GRCm39) using BWA-MEM with default parameters ^12^. The resulting SAM files were converted to BAM format and sorted using SAMtools ^15,19^. PCR duplicates were marked using GATK MarkDuplicatesSpark ^14^. Read groups were added to the BAM files using GATK AddOrReplaceReadGroups. BQSR was performed using GATK BaseRecalibrator with known SNPs and indels from the Mouse Genome Project (MGP) ^20,21^. The recalibrated BAM files were then used for variant calling. Variants were called using GATK HaplotypeCaller in GVCF mode. The GVCF files for each sample were combined using GATK CombineGVCFs, followed by joint genotyping with GATK GenotypeGVCFs to produce the final VCF files. Variants were annotated using Variant Effect Predictor (VEP, release 107)^22^ with the mouse GRCm39 cache. The annotations included predicted functional impacts (HIGH, MODERATE, LOW, MODIFIER) and other relevant information. High-confidence variants were filtered based on QD (Quality by Depth) and other relevant metrics using BCFtools (v 1.16) ^15^. Specifically, variants with QD < 2 were removed. Variants with HIGH and MODERATE impacts were retained for further analysis. Variants located in the specific region chr7:58878550-60099925 were extracted using bcftools.

### CHANGE-seq

Genomic DNA extracted from 3 unedited iPSC lines from healthy donor using Gentra Puregene kit (Qiagen) according to the manufacturer’s recommendation. CHANGE-seq was performed as previously described ^23^. In short, a custom Tn5-transposome used to tagment the purified genomic DNA to an average length of 500-800 bp, and was gap repaired with HiFi HotStart Uracil+ Ready Mix (Kapa Biosystems). USER enzyme (NEB) and T4 polyncucleotide was then used to treat the gap-repaired DNA. DNA circularization was performed using T4 DNA ligase (NEB) and residual traces of linear DNA was removed by a cocktail of exonucleases. *In vitro* cleavage reactions were performed on the circularized DNA using *Sp*Cas9 protein (NEB) and gRNA. Cleaved products were incubated with proteinase K (NEB), A-tailed, hairpin adapter (NEB) ligated, and treated with USER enzyme (NEB). Products were amplified by PCR with the Kapa HiFi polymerase (Kapa Biosystems), and libraries were quantified by qPCR (Kapa Biosystems). Libraries were pooled, loaded and sequenced on the NextSeq2000 with the following cycles 151-8-8-151 according to the manufacturer’s instructions. CHANGE-seq data analyses were performed as reported previously^23^.

### Twist Hybridization Analysis

A custom-designed NGS enrichment panel consisting of 3,738 probes targeting 4,421 CHANGE-seq predicted on- and off-target regions (Design ID TE-99523833, Twist Bioscience) was synthesized. Three iPSC lines used for CHANGE-seq nomination, one non-biological relevant iPSC line, and 4 induced neuron lines (iNeurons) from these 4 iPSCs underwent targeted enrichment. Hybridization capture enrichment was conducted at the Yale Center for Genome Analysis (YCGA) using standardized protocol, followed by sequencing on the Illumina platform to achieve approximately 32 million reads per sample. Raw FASTQ data were processed using following 3 pipelines:

**1.** Quality assessment of raw sequencing reads was conducted using fastp^16^. Adapter trimming and quality filtering were performed with Trim Galore to ensure high-quality reads. Filtered reads were aligned to the human reference genome (hg38) using BWA-MEM with default parameters^12^. Alignment outputs (SAM format) were converted to BAM files, sorted, and indexed using SAMtools ^15,19^. Reads were subsequently narrowed to probe-targeted regions. PCR duplicates were marked using GATK MarkDuplicatesSpark^14^, and read groups were added using GATK AddOrReplaceReadGroups. Variants were then called from the deduplicated BAM file using GATK HaplotypeCaller.
**2.** Paired-end reads were trimmed using Cutadapt ^24^ with paramaters -m 20 -O 5 -a AGATCGGAAGAGC -A AGATCGGAAGAGC. Trimmed reads were then merged using FLASH^25^. Both merged and unmerged FASTQ files were concatenated and aligned to the hg38 reference genome using BWA-MEM as described above. On- and off-target indel frequencies were quantified using CRISPResso2 (version 2.3.2) ^26^ with parameters -r hg38.fa --quantification_window_size 1 --quantification_window_center -3 --base_editor_output -- min_reads_to_use_region 50 -b $bam --exclude_bp_from_right 0 --exclude_bp_from_left 0 --plot_window_size 12. Regions with fewer than 1000 total reads were filtered out.
**3.** For the CRISPResso-specific pipeline, FASTQ files from each STEP-RNP treated and corresponding untreated control sample were individually analyzed using CRISPResso2 (version 2.3.2) via CRISPRessoPooled Amplicons Mode, including reads merging and trimming using fastp, and aligning to amplicon sequences with bowtie2, and results were subsequently compared using CRISPRessoCompare mode of CRISPResso2 with default parameters.

### PacBio HiFi Sequence and Analysis

The same iPSC and iNeuron lines described above were subjected to PacBio HiFi sequencing. High molecular weight (HMW) genomic DNA was isolated using the PacBio Nanobind HMW DNA Extraction Kit (Nanobind PanDNA 103-260-000) according to the manufacturer’s instructions, optimized for SMRT library preparation. DNA libraries were sequenced using one SMRTcell per sample on PacBio Revio system at the Yale Center for Genome Analysis (YCGA). Resulting HiFi reads were subsequently analyzed. Off-target editing events were identified using the HiFi-somatic-WDL pipeline with default parameters, capturing small variants (single nucleotide variants [SNVs] and insertions/deletions [INDELs]), structural variants (SVs), copy number variants, and methylation differences by comparing each STEP-RNP treated sample to its matched untreated control. For on-target analysis, reads were specifically narrowed to the genomic region chr15:25193313–25439024 using SAMtools. Targeted analyses included read alignment with pbmm2, variant calling for SNVs and small INDELs using DeepVariant^27^, SV detection using pbsv, and DNA methylation probability assessment at CpG sites using pb-CpG-tools. Variants were subsequently annotated: small variants were annotated using slivar^28^, and SV annotations were performed using svpack.

### Data Visualization

Visualization was performed using ggplot2 (version 3.3.2) in R (version 4.2.2) for plotting gene/genome structure, single nucleotide variant, genome structural variant, gene expression and heatmaps.

### Data and Statistical analysis

For groups or conditions with more than two categories, statistical analyses were performed using Bonferroni-corrected pair-wise comparisons and Dunnett’s post-hoc tests, unless otherwise specified. In all cases, a p-value of less than 0.05 was considered significant. However, for the immune and inflammatory protein-based panel, multiple t-tests were employed using the False Discovery Rate (FDR) approach, specifically the two-stage step-up method of Benjamini, Krieger, and Yekutieli, with a desired FDR (Q) set at 1%.

## References

1 Musunuru, K. et al. Patient-Specific In Vivo Gene Editing to Treat a Rare Genetic Disease. N Engl J Med 392, 2235–2243 (2025). 10.1056/NEJMoa2504747

2 Frangoul, H. et al. Exagamglogene Autotemcel for Severe Sickle Cell Disease. N Engl J Med 390, 1649–1662 (2024). 10.1056/NEJMoa2309676

3 Fontana, M. et al. CRISPR-Cas9 Gene Editing with Nexiguran Ziclumeran for ATTR Cardiomyopathy. N Engl J Med 391, 2231–2241 (2024). 10.1056/NEJMoa2412309

4 Cohn, D. M. et al. CRISPR-Based Therapy for Hereditary Angioedema. N Engl J Med 392, 458–467 (2025). 10.1056/NEJMoa2405734

5 Chen, G. et al. A biodegradable nanocapsule delivers a Cas9 ribonucleoprotein complex for in vivo genome editing. Nat Nanotechnol 14, 974–980 (2019). 10.1038/s41565-019-0539-2

6 Foss, D. V. et al. Peptide-mediated delivery of CRISPR enzymes for the efficient editing of primary human lymphocytes. Nat Biomed Eng 7, 647–660 (2023). 10.1038/s41551-023-01032-2

7 Kulhankova, K., et al. Shuttle peptide delivers base editor RNPs to rhesus monkey airway epithelial cells in vivo. Nat Commun 14, 8051 (2023). 10.1038/s41467-023-43904-w

8 Staahl, B. T. et al. Efficient genome editing in the mouse brain by local delivery of engineered Cas9 ribonucleoprotein complexes. Nat Biotechnol 35, 431–434 (2017). 10.1038/nbt.3806

9 Lee, K. et al. Nanoparticle delivery of Cas9 ribonucleoprotein and donor DNA in vivo induces homology-directed DNA repair. Nat Biomed Eng 1, 889–901 (2017). 10.1038/s41551-017-0137-2

10 Saha, K. et al. The NIH Somatic Cell Genome Editing program. Nature 592, 195–204 (2021). 10.1038/s41586-021-03191-1

11 Tsuchida, C. A., Wasko, K. M., Hamilton, J. R. & Doudna, J. A. Targeted nonviral delivery of genome editors in vivo. Proc Natl Acad Sci U S A 121, e2307796121 (2024). 10.1073/pnas.2307796121

12 Chen, K. et al. Engineering self-deliverable ribonucleoproteins for genome editing in the brain. Nat Commun 15, 1727 (2024). 10.1038/s41467-024-45998-2

13 Stahl, E. C. et al. Genome editing in the mouse brain with minimally immunogenic Cas9 RNPs. Mol Ther 31, 2422–2438 (2023). 10.1016/j.ymthe.2023.06.019

14 Gao, J. et al. Gene therapy for CNS disorders: modalities, delivery and translational challenges. Nat Rev Neurosci 25, 553–572 (2024). 10.1038/s41583-024-00829-7

15 Goertsen, D. et al. AAV capsid variants with brain-wide transgene expression and decreased liver targeting after intravenous delivery in mouse and marmoset. Nat Neurosci 25, 106–115 (2022). 10.1038/s41593-021-00969-4

16 Challis, R. C. et al. Systemic AAV vectors for widespread and targeted gene delivery in rodents. Nat Protoc 14, 379–414 (2019). 10.1038/s41596-018-0097-3

17 Kondratov, O. et al. A comprehensive study of a 29-capsid AAV library in a non-human primate central nervous system. Mol Ther 29, 2806–2820 (2021). 10.1016/j.ymthe.2021.07.010

18 Hordeaux, J. et al. The GPI-Linked Protein LY6A Drives AAV-PHP.B Transport across the Blood-Brain Barrier. Mol Ther 27, 912–921 (2019). 10.1016/j.ymthe.2019.02.013

19 Li, J. et al. A high-fidelity RNA-targeting Cas13 restores paternal Ube3a expression and improves motor functions in Angelman syndrome mice. Mol Ther 31, 2286–2295 (2023). 10.1016/j.ymthe.2023.02.015

20 Huang, Q. et al. An AAV capsid reprogrammed to bind human transferrin receptor mediates brain-wide gene delivery. Science 384, 1220–1227 (2024). 10.1126/science.adm8386

21 Li, W. K. et al. Whole-brain in vivo base editing reverses behavioral changes in Mef2c-mutant mice. Nat Neurosci 27, 116–128 (2024). 10.1038/s41593-023-01499-x

22 Greig, J. A. et al. Integrated vector genomes may contribute to long-term expression in primate liver after AAV administration. Nat Biotechnol (2023). 10.1038/s41587-023-01974-7

23 Hordeaux, J. et al. MicroRNA-mediated inhibition of transgene expression reduces dorsal root ganglion toxicity by AAV vectors in primates. Sci Transl Med 12 (2020). 10.1126/scitranslmed.aba9188

24 Chowdary, P. et al. Phase 1-2 Trial of AAVS3 Gene Therapy in Patients with Hemophilia B. N Engl J Med 387, 237–247 (2022). 10.1056/NEJMoa2119913

25 Arjomandnejad, M., Dasgupta, I., Flotte, T. R. & Keeler, A. M. Immunogenicity of Recombinant Adeno-Associated Virus (AAV) Vectors for Gene Transfer. BioDrugs 37, 311–329 (2023). 10.1007/s40259-023-00585-7

26 Vrellaku, B. et al. A systematic review of immunosuppressive protocols used in AAV gene therapy for monogenic disorders. Mol Ther 32, 3220–3259 (2024). 10.1016/j.ymthe.2024.07.016

27 Shirley, J. L., de Jong, Y. P., Terhorst, C. & Herzog, R. W. Immune Responses to Viral Gene Therapy Vectors. Mol Ther 28, 709–722 (2020). 10.1016/j.ymthe.2020.01.001

28 Zou, Y. et al. Blood-brain barrier-penetrating single CRISPR-Cas9 nanocapsules for effective and safe glioblastoma gene therapy. Sci Adv 8, eabm8011 (2022). 10.1126/sciadv.abm8011

29 Shergalis, A., Bankhead, A., 3rd, Luesakul, U., Muangsin, N. & Neamati, N. Current Challenges and Opportunities in Treating Glioblastoma. Pharmacol Rev 70, 412–445 (2018). 10.1124/pr.117.014944

30 Thorne, R. G. & Nicholson, C. In vivo diffusion analysis with quantum dots and dextrans predicts the width of brain extracellular space. Proc Natl Acad Sci U S A 103, 5567–5572 (2006). 0509425103 [pii] 10.1073/pnas.0509425103

31 Chen, Z. et al. Targeted Delivery of CRISPR/Cas9-Mediated Cancer Gene Therapy via Liposome-Templated Hydrogel Nanoparticles. Adv Funct Mater 27 (2017). 10.1002/adfm.201703036

32 Jiang, Y., Lev-Lehman, E., Bressler, J., Tsai, T. F. & Beaudet, A. L. Genetics of Angelman syndrome. Am J Hum Genet 65, 1–6. (1999).

33 Buiting, K., Williams, C. & Horsthemke, B. Angelman syndrome - insights into a rare neurogenetic disorder. Nat Rev Neurol 12, 584–593 (2016). 10.1038/nrneurol.2016.133

34 Albrecht, U. et al. Imprinted expression of the murine Angelman syndrome gene, Ube3a, in hippocampal and Purkinje neurons. Nat Genet 17, 75–78. (1997).

35 Meng, L., Person, R. E. & Beaudet, A. L. Ube3a-ATS is an atypical RNA polymerase II transcript that represses the paternal expression of Ube3a. Hum Mol Genet 21, 3001–3012 (2012). dds130 [pii] 10.1093/hmg/dds130

36 Meng, L. et al. Truncation of Ube3a-ATS unsilences paternal Ube3a and ameliorates behavioral defects in the Angelman syndrome mouse model. PLoS Genet 9, e1004039 (2013). 10.1371/journal.pgen.1004039

37 Meng, L. et al. Towards a therapy for Angelman syndrome by targeting a long non-coding RNA. Nature 518, 409–412 (2015). 10.1038/nature13975

38 Bailus, B. J. et al. Protein Delivery of an Artificial Transcription Factor Restores Widespread Ube3a Expression in an Angelman Syndrome Mouse Brain. Mol Ther 24, 548–555 (2016). 10.1038/mt.2015.236

39 Huang, H. S. et al. Topoisomerase inhibitors unsilence the dormant allele of Ube3a in neurons. Nature 481, 185–189 (2011). 10.1038/nature10726

40 Schmid, R. S. et al. CRISPR/Cas9 directed to the Ube3a antisense transcript improves Angelman syndrome phenotype in mice. J Clin Invest 131 (2021). 10.1172/jci142574

41 Wolter, J. M. et al. Cas9 gene therapy for Angelman syndrome traps Ube3a-ATS long non-coding RNA. Nature 587, 281–284 (2020). 10.1038/s41586-020-2835-2

42 Berry-Kravis E, O. C., Richter H, Nolan M, Kakkis E, Brandabur M, Dindot S, Panagoulias J, Stromatt S. Early Results from a Five Patient Cohort Treated with GTX-102, An Antisense Oligonucleotide Designed to Activate the Paternal UBE3A Gene in Angelman Syndrome. Early Results from a Five Patient Cohort Treated with GTX-102, An Antisense Oligonucleotide Designed to Activate the Paternal UBE3A Gene in Angelman Syndrome. Ann Neurol. (2021).

43 Hipp, J. F. et al. The UBE3A-ATS antisense oligonucleotide rugonersen in children with Angelman syndrome: a phase 1 trial. Nat Med 31, 2936–2945 (2025). 10.1038/s41591-025-03784-7

44 Cox, D. B. T. et al. RNA editing with CRISPR-Cas13. Science 358, 1019–1027 (2017). 10.1126/science.aaq0180

45 Tabebordbar, M. et al. In vivo gene editing in dystrophic mouse muscle and muscle stem cells. Science 351, 407–411 (2016). 10.1126/science.aad5177

46 Madisen, L. et al. A robust and high-throughput Cre reporting and characterization system for the whole mouse brain. Nat Neurosci 13, 133–140 (2010). 10.1038/nn.2467

47 Dindot, S. V., Antalffy, B. A., Bhattacharjee, M. B. & Beaudet, A. L. The Angelman syndrome ubiquitin ligase localizes to the synapse and nucleus, and maternal deficiency results in abnormal dendritic spine morphology. Hum Mol Genet 17, 111–118 (2008). 10.1093/hmg/ddm288

48 Yamasaki, K. et al. Neurons but not glial cells show reciprocal imprinting of sense and antisense transcripts of Ube3a. Hum Mol Genet 12, 837–847 (2003). 10.1093/hmg/ddg106

49 Judson, M. C., Sosa-Pagan, J. O., Del Cid, W. A., Han, J. E. & Philpot, B. D. Allelic specificity of Ube3a expression in the mouse brain during postnatal development. J Comp Neurol 522, 1874–1896 (2014). 10.1002/cne.23507

50 Schmid, R. S. et al. CRISPR/Cas9 directed to the Ube3a antisense transcript improves Angelman syndrome phenotype in mice. J Clin Invest (2021). 10.1172/JCI142574

51 Lu, X. & Jiang, Y. Inerathecal Injection of Newborn Mouse for Genome Editing and Drug Delivery. JOVE (2023).

52 Bieth, E. et al. Highly restricted deletion of the SNORD116 region is implicated in Prader-Willi Syndrome. Eur J Hum Genet 23, 252–255 (2015). 10.1038/ejhg.2014.103

53 Sahoo, T. et al. Prader-Willi phenotype caused by paternal deficiency for the HBII-85 C/D box small nucleolar RNA cluster. Nat Genet 40, 719–721 (2008).

54 Dindot, S. V., Antalffy, B. A., Bhattacharjee, M. B. & Beaudet, A. L. The Angelman syndrome ubiquitin ligase localizes to the synapse and nucleus, and maternal deficiency results in abnormal dendritic spine morphology. Human Molecular Genetics 17, 111–118 (2007). 10.1093/hmg/ddm288

55 Jiang, Y. H. et al. Mutation of the Angelman ubiquitin ligase in mice causes increased cytoplasmic p53 and deficits of contextual learning and long-term potentiation. Neuron 21, 799–811 (1998).

56 Sun, A. X. et al. Potassium channel dysfunction in human neuronal models of Angelman syndrome. Science 366, 1486–1492 (2019). 10.1126/science.aav5386

57 Keute, M. et al. Angelman syndrome genotypes manifest varying degrees of clinical severity and developmental impairment. Mol Psychiatry 26, 3625–3633 (2021). 10.1038/s41380-020-0858-6

58 Egawa, K. et al. Flurothyl-induced seizure paradigm revealed higher seizure susceptibility in middle-aged Angelman syndrome mouse model. Brain Dev 43, 515–520 (2021). 10.1016/j.braindev.2020.12.011

59 Gu, B. et al. Ube3a reinstatement mitigates epileptogenesis in Angelman syndrome model mice. J Clin Invest 129, 163–168 (2019). 10.1172/jci120816

60 Williams, C. A. et al. Angelman syndrome 2005: updated consensus for diagnostic criteria. Am J Med Genet A 140, 413–418 (2006). 10.1002/ajmg.a.31074

61 Dindot, S. V. et al. An ASO therapy for Angelman syndrome that targets an evolutionarily conserved region at the start of the UBE3A-AS transcript. Sci Transl Med 15, eabf4077 (2023). 10.1126/scitranslmed.abf4077

62 Zhang, Y. et al. Rapid single-step induction of functional neurons from human pluripotent stem cells. Neuron 78, 785–798 (2013). 10.1016/j.neuron.2013.05.029

63 Doran, A. G. et al. Deep genome sequencing and variation analysis of 13 inbred mouse strains defines candidate phenotypic alleles, private variation and homozygous truncating mutations. Genome Biology 17, 167 (2016). 10.1186/s13059-016-1024-y

64 Keane, T. M. et al. Mouse genomic variation and its effect on phenotypes and gene regulation. Nature 477, 289–294 (2011). 10.1038/nature10413

65 Bae, S., Park, J. & Kim, J.-S. Cas-OFFinder: a fast and versatile algorithm that searches for potential off-target sites of Cas9 RNA-guided endonucleases. Bioinformatics 30, 1473–1475 (2014). 10.1093/bioinformatics/btu048

66 Lazzarotto, C. R. et al. CHANGE-seq reveals genetic and epigenetic effects on CRISPR-Cas9 genome-wide activity. Nat Biotechnol 38, 1317–1327 (2020). 10.1038/s41587-020-0555-7

67 Hiragi, T., Ikegaya, Y. & Koyama, R. Microglia after Seizures and in Epilepsy. Cells 7 (2018). 10.3390/cells7040026

68 Le Fevre, A. et al. Atypical Angelman syndrome due to a mosaic imprinting defect: Case reports and review of the literature. Am J Med Genet A 173, 753–757 (2017). 10.1002/ajmg.a.38072

69 Nazlican, H. et al. Somatic mosaicism in patients with Angelman syndrome and an imprinting defect. Hum Mol Genet 13, 2547–2555 (2004). 10.1093/hmg/ddh296

70 Ramadoss, G. N. et al. Neuronal DNA repair reveals strategies to influence CRISPR editing outcomes. bioRxiv (2024). 10.1101/2024.06.25.600517

## References

1 Chen, K. et al. Engineering self-deliverable ribonucleoproteins for genome editing in the brain. Nat Commun 15, 1727 (2024). 10.1038/s41467-024-45998-2

2 Zhang, Y. et al. Rapid single-step induction of functional neurons from human pluripotent stem cells. Neuron 78, 785–798 (2013). 10.1016/j.neuron.2013.05.029

3 Xiang, Y. et al. Generation and Fusion of Human Cortical and Medial Ganglionic Eminence Brain Organoids. Curr Protoc Stem Cell Biol 47 (2018). 10.1002/cpsc.61

4 Madisen, L. et al. A robust and high-throughput Cre reporting and characterization system for the whole mouse brain. Nat Neurosci 13, 133–140 (2010). 10.1038/nn.2467

5 Dindot, S. V., Antalffy, B. A., Bhattacharjee, M. B. & Beaudet, A. L. The Angelman syndrome ubiquitin ligase localizes to the synapse and nucleus, and maternal deficiency results in abnormal dendritic spine morphology. Hum Mol Genet 17, 111–118 (2008). 10.1093/hmg/ddm288

6 Jiang, Y. H. et al. Mutation of the Angelman ubiquitin ligase in mice causes increased cytoplasmic p53 and deficits of contextual learning and long-term potentiation. Neuron 21, 799–811 (1998).

7 Lu, X. & Jiang, Y.-h. Intrathecal Injection of Newborn Mouse for Genome Editing and Drug Delivery. JoVE, e65761 doi:10.3791/65761

8 Kim, J. Y., Grunke, S. D., Levites, Y., Golde, T. E. & Jankowsky, J. L. Intracerebroventricular viral injection of the neonatal mouse brain for persistent and widespread neuronal transduction. J Vis Exp, 51863 (2014). 10.3791/51863

9 Gombash Lampe, S. E., Kaspar, B. K. & Foust, K. D. Intravenous injections in neonatal mice. J Vis Exp, e52037 (2014). 10.3791/52037

10 Egawa, K. et al. Flurothyl-induced seizure paradigm revealed higher seizure susceptibility in middle-aged Angelman syndrome mouse model. Brain Dev 43, 515–520 (2021). 10.1016/j.braindev.2020.12.011

11 Martin, M. Cutadapt removes adapter sequences from high-throughput sequencing reads. EMBnet.journal 17, 3 (2011). 10.14806/ej.17.1.200

12 Jung, Y. & Han, D. BWA-MEME: BWA-MEM emulated with a machine learning approach. Bioinformatics 38, 2404–2413 (2022). 10.1093/bioinformatics/btac137

13 toolkit, P. Picard toolkit, <https://broadinstitute.github.io/picard/> (2019).

14 McKenna, A. et al. The Genome Analysis Toolkit: a MapReduce framework for analyzing next-generation DNA sequencing data. Genome Res 20, 1297–1303 (2010). 10.1101/gr.107524.110

15 Danecek, P. et al. Twelve years of SAMtools and BCFtools. GigaScience 10 (2021). 10.1093/gigascience/giab008

16 Chen, S., Zhou, Y., Chen, Y. & Gu, J. fastp: an ultra-fast all-in-one FASTQ preprocessor. Bioinformatics 34, i884–i890 (2018). 10.1093/bioinformatics/bty560

17 Dobin, A. et al. STAR: ultrafast universal RNA-seq aligner. Bioinformatics 29, 15–21 (2013). 10.1093/bioinformatics/bts635

18 Liao, Y., Smyth, G. K. & Shi, W. featureCounts: an efficient general purpose program for assigning sequence reads to genomic features. Bioinformatics 30, 923–930 (2014). 10.1093/bioinformatics/btt656

19 Li, H. et al. The Sequence Alignment/Map format and SAMtools. Bioinformatics 25, 2078–2079 (2009). 10.1093/bioinformatics/btp352

20 Doran, A. G. et al. Deep genome sequencing and variation analysis of 13 inbred mouse strains defines candidate phenotypic alleles, private variation and homozygous truncating mutations. Genome Biology 17, 167 (2016). 10.1186/s13059-016-1024-y

21 Keane, T. M. et al. Mouse genomic variation and its effect on phenotypes and gene regulation. Nature 477, 289–294 (2011). 10.1038/nature10413

22 McLaren, W. et al. The Ensembl Variant Effect Predictor. Genome Biol 17, 122 (2016). 10.1186/s13059-016-0974-4

23 Lazzarotto, C. R. et al. CHANGE-seq reveals genetic and epigenetic effects on CRISPR-Cas9 genome-wide activity. Nat Biotechnol 38, 1317–1327 (2020). 10.1038/s41587-020-0555-7

24 Martin, M. Cutadapt removes adapter sequences from high-throughput sequencing reads. 2011 17, 3 (2011). 10.14806/ej.17.1.200

25 Magoč, T. & Salzberg, S. L. FLASH: fast length adjustment of short reads to improve genome assemblies. Bioinformatics 27, 2957–2963 (2011). 10.1093/bioinformatics/btr507

26 Clement, K. et al. CRISPResso2 provides accurate and rapid genome editing sequence analysis. Nat Biotechnol 37, 224–226 (2019). 10.1038/s41587-019-0032-3

27 Poplin, R. et al. A universal SNP and small-indel variant caller using deep neural networks. Nature Biotechnology 36, 983–987 (2018). 10.1038/nbt.4235

28 Pedersen, B. S. et al. Effective variant filtering and expected candidate variant yield in studies of rare human disease. npj Genomic Medicine 6, 60 (2021). 10.1038/s41525-021-00227-3

