## Supplementary material for "Brain-wide Genome Editing via STEP-RNPs for Treatment of Angelman Syndrome": Extend Fig

### Extended Data 1

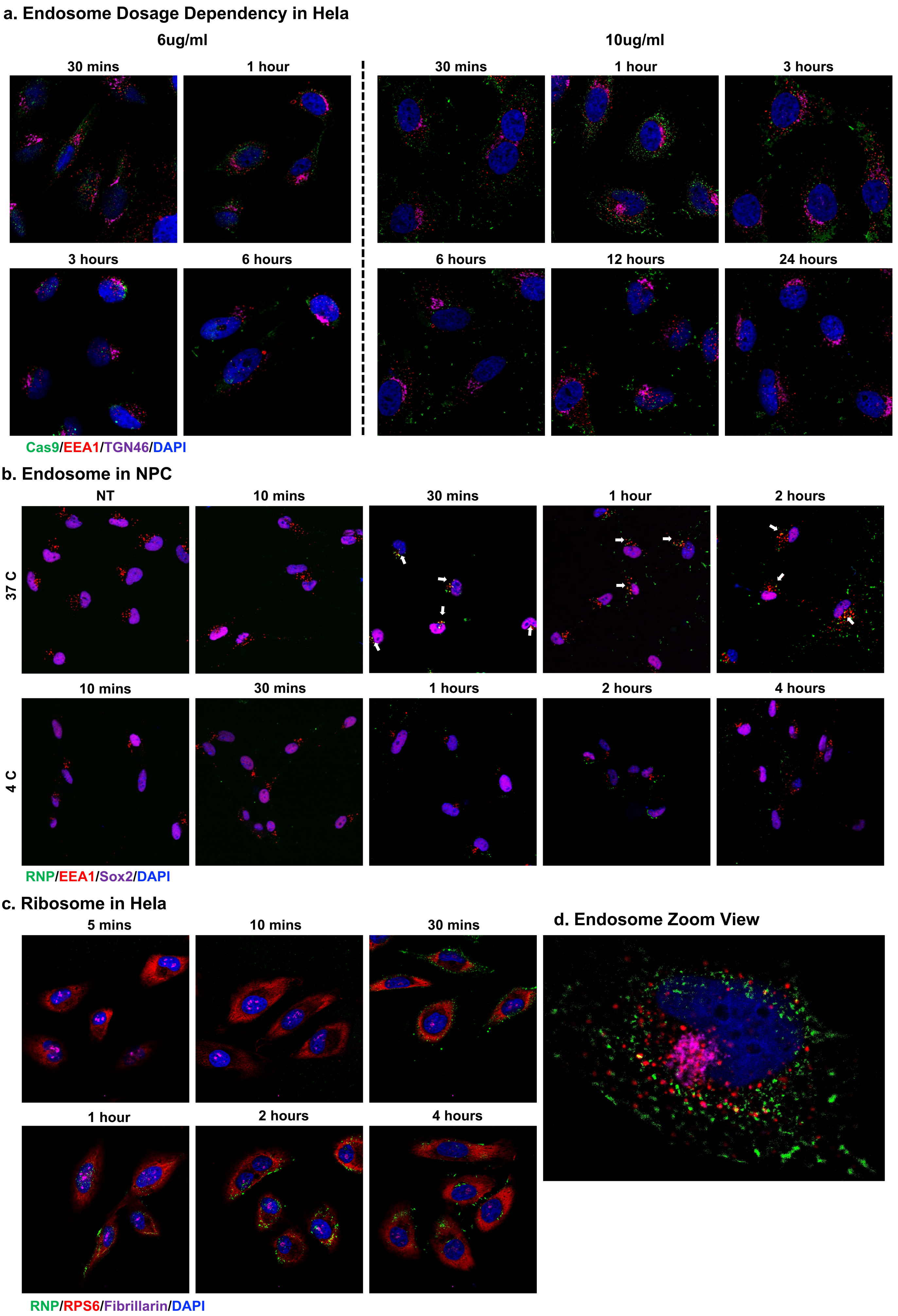

**Extended Data 1**

**Intracellular localization of STEP-RNP over time following uptake in HeLa cells (a,c,d) and neural progenitor cell (b).**

**(a)** HeLa cells were treated with STEP-RNP at Cas9 concentrations of 6 μg/mL (left) and 10 μg/mL (right) and fixed at the indicated time points (30 mins, 1 hour, 3 hours, 6 hours, 12 hours, and 24 hours) post-treatment. Immunofluorescence staining was performed for Cas9 (green), early endosome marker EEA1 (red), trans-Golgi network marker TGN46 (magenta), and nuclei (DAPI, blue). Images show the dynamic intracellular trafficking of STEP-RNP, with early colocalization in endosomes (EEA1) followed by redistribution over time.

**(b)** Temperature-dependent endosomal localization of RNP in neural progenitor cells (NPCs). NPCs were treated with STEP-RNP (green) and fixed at the indicated time points after incubation at either 37°C (top) or 4°C (bottom). Immunofluorescence staining was performed for the early endosome marker EEA1 (red), the neural progenitor marker Sox2 (magenta), and nuclei (DAPI, blue). At 37°C, RNP progressively colocalizes with EEA1-positive endosomes started within 30 minutes (arrows), indicating active endocytosis over time. In contrast, at 4°C, endosomal uptake is reduced, suggesting temperature-dependent internalization.

**(c)** Ribosome intake of STEP-RNP in Hela cells. STEP-RNP co-strained with mature ribosome marker RPS6 (red), nucleolar ribosome biogenesis marker Fibrillarin (magenta), and nuclei (DAPI, blue). Throughout the time course, STEP-RNP exhibited limited colocalization with RPS6, but no Fibrillarin. This is to clarify the endocytosis pathway, tested in Hela cells.

**(d)** Zoomed in view of co-localization of STEP-RNP and endosome, 1 hour after 10ug/ml of STEP-RNP application.

#### Extended Data 2

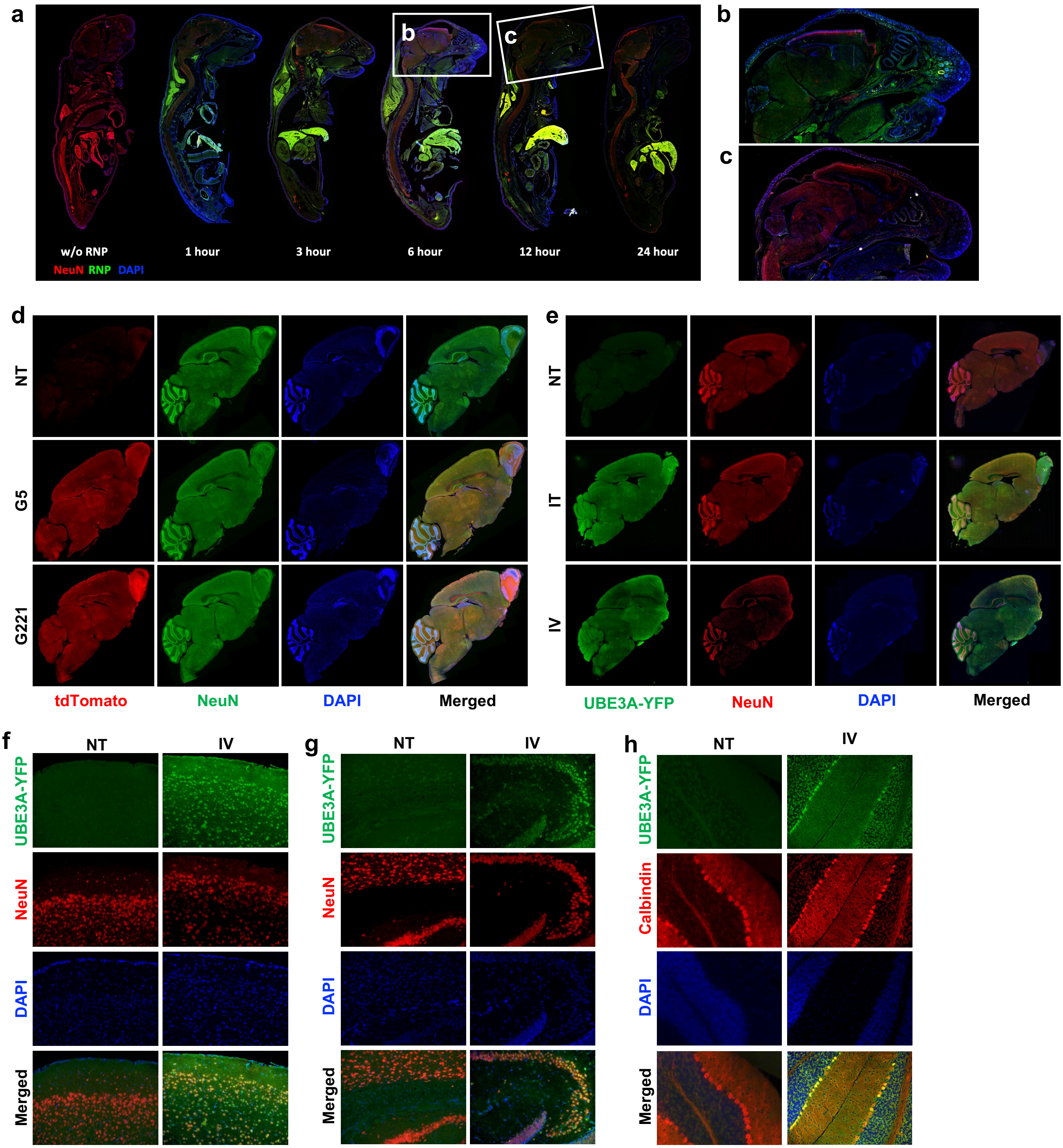

**Extended Data 2**

**Biodistribution of STEP-RNP delivered via IV (a-c), brain-wide gene editing of STEP-RNP delivered via IT in Ai9 reporter mouse model (d) and IV and IT in Ube3a-YFP reporter mice (e-f)**

**(a-c)** Biodistribution of STEP-RNP in mouse over time (1, 3, 6, 12, and 24 hours) after IV at P1 day (10 µg/µL, 10 µL/mouse). STEP-RNP was detected by immunostaining of the His-tag fused with G5 Cas9. STEP-RNP was detectable in the brain within 1 hour following administration and became undetectable in the brain by 12 hours (n = 6 per time point, scale bar = 5 mm).

**(d)** Brain-wide gene editing of STEP-RNP delivered in Ai9 reporter mouse model. Brain-wide gene editing effect of STEP-RNP synthesized with two different versions of Cas9 (G5 and G221) delivered to Ai9 reporter mice on P1 via IT injection compared to the NT group. G221 differs from G5 with a modified version of NLS. The editing effect was assessed 30 days after IT administration. Both variants of Cas9 demonstrated high and comparable brain-wide genome editing efficacy but noted that G221 Cas9 displayed more efficient editing in the olfactory bulb compared to G5 Cas9 (n=6 per group).

**(e)** Brain-wide gene editing effect of STEP-RNP delivered via IV and IT, 30 days after injection in P1 in *Ube3am*$^{+/p\text{-}YFP}$ mice. Overall, STEP-RNP achieved better brain-wide editing through IT delivery than IV.

**(f)** Efficient gene editing by STEP-RNP delivered via IV in NeuN+ neurons of the prefrontal cortex (NT, n=12; IV, n=6).

**(g)** Efficient gene editing by STEP-RNP delivered via IV in NeuN+ neurons of the hippocampus. (NT, n=12; IV, n=6).

**(h)** Efficient gene editing efficacy of STEP-RNP delivered via IV in calbindin+ Purkinje cells in the cerebellum (NT, n=12; IV, n=6).

### Extended Data 3

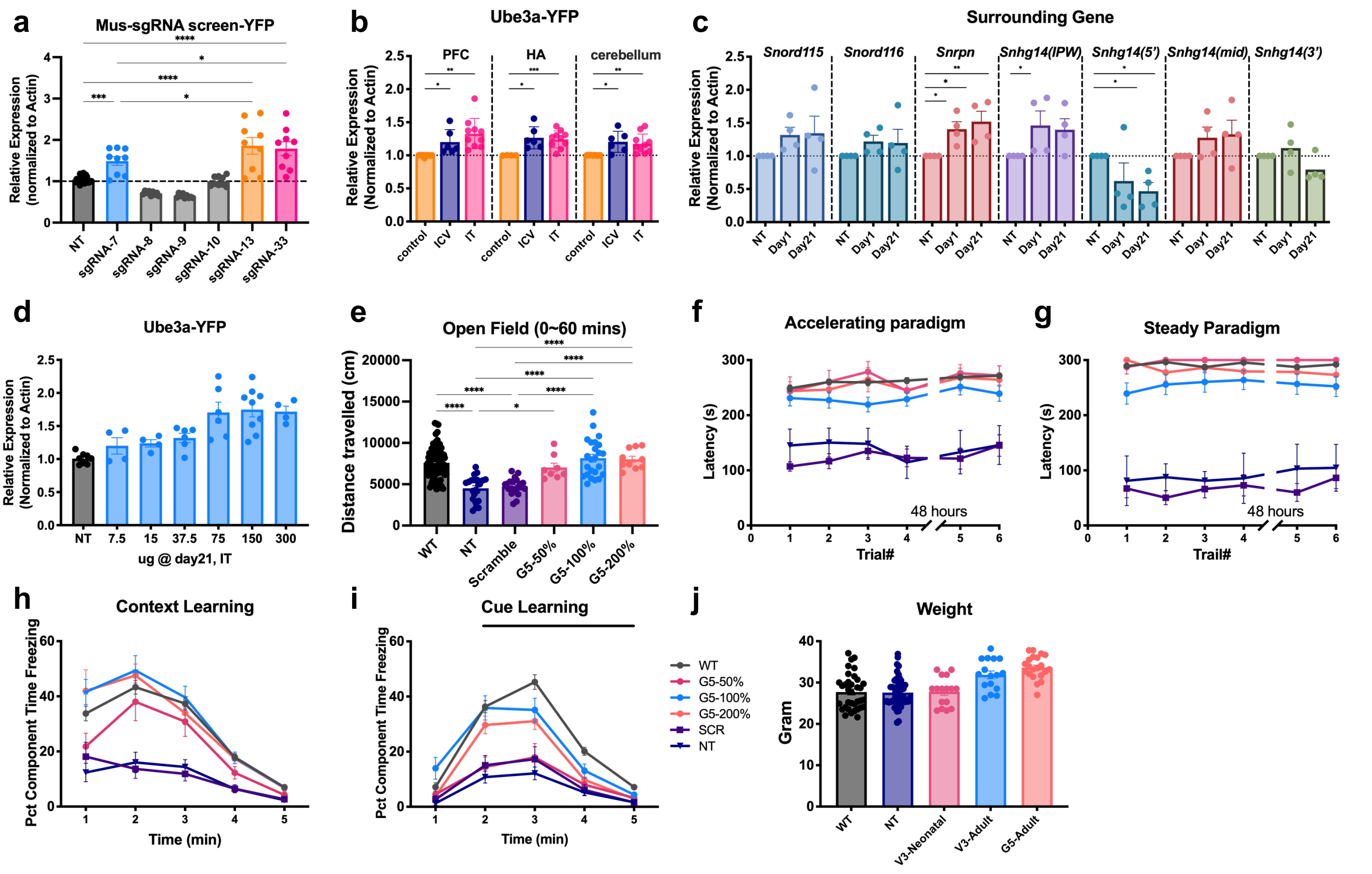


**Extended Data 3**

**sgRNA screening and Dosage response of STEP-RNP treatment in AS reporter and AS mouse models.**

**(a)** Screening of selected sgRNAs revealed that sgRNA33 is more effective than other sgRNAs (sgRNA-7, sgRNA-8, sgRNA-9 and sgRNA-10).

**(b)** The editing efficiency of ICV and IT deliveries at P1 are comparable in Ube3a-YFP reporter mice assessed 30 days after injection. (**: p<0.01; ***: p<0.001)

**(c)** Relative expression of genes surrounding *Ube3a-ATS* as assessed by qRT-PCR. Notably, there were no significant changes in the expression levels of *Snord115* and *Snord116* in the prefrontal cortex of mice treated with STEP-RNP when compared to untreated controls. A consistent decrease in the expression of *Snhg14* at the 5' end paralleled the observations in the qRT-PCR of *Ube3a-ATS* (n=4 for each group, *p<0.05; ** p<0.01).

**(d)** Different dosage of STEP-RNP quantified by the amount of Cas9 protein, from 7.5 to 300 ug in 15 ul volume, was administrated to Ube3a-YFP reporter mice via IT route. The 150 ug is defined as equivalent to the 100% dosage used in this study. RT-qPCR showed the 75 ug is the minimal effective dosage for the reactivation UBE3A-YFP RNA level.

**(e-i)** Different dosage of STEP-RNP in AS mouse models. Based on the assessment of multiple functional domains of behaviors (locomotor, motor balance, cognitive), 50% of STEP-RNP is the minimum effective dose for open field, rotarod, and contextual fear conditioning but 100% is minimum effective dosage for the cured conditioning.

**(j)** Body weight of AS mice at 5-6 months after treated with STEP-RNP in different groups is comparable with nontreated and WT mice.

### Extended Data 4

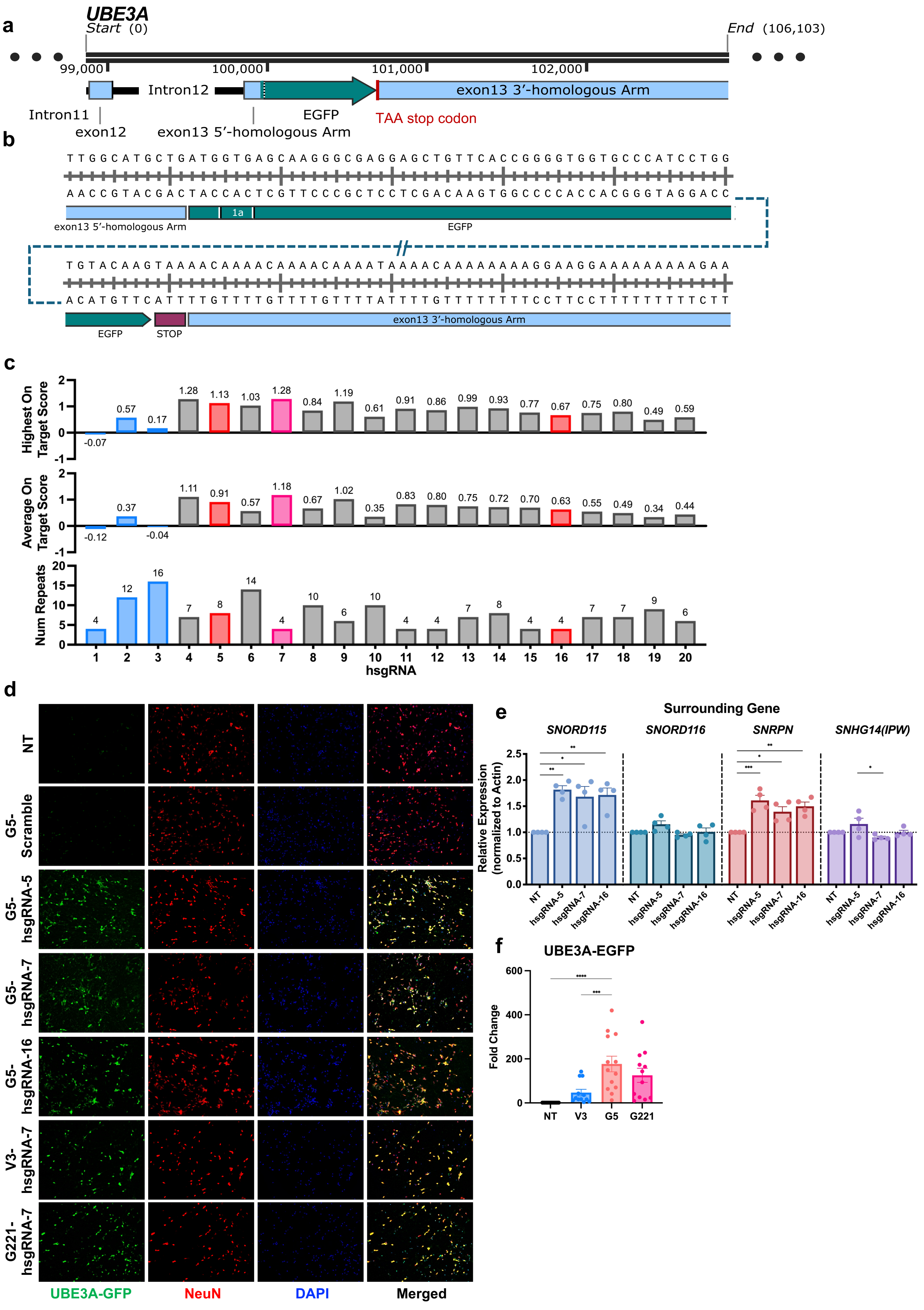

**Extended Data 4**

**(a)** Paternal UBE3A-EGFP fusion hIPSCs line design. *UBE3A* and *EGFP* fusion gene structure.

**(b)** *UBE3A* and *EGFP* fusion gene sequence. EGFP was fused in-frame to UBE3A after disruption of STOP codon of UBE3A

**(c)** In silico analysis of sgRNAs candidates targeting UBE3A-ATS. The design of the human-specific sgRNA (hsgRNA) targets the cross species conserved UBE3A-ATS genomic region. hsgRNA candidates were selected according to the target score and number of targets from in silico analysis. Blue bars represent previously published hsgRNAs. The pink hsgRNA-7 has the highest on-target scores, red bars represent other hsgRNAs tested in following experiments, and grey bars represent the remaining hsgRNAs.

**(d)** Representative images of editing effect of 20 hsgRNA candidates in AS-hIPSCs-iNeurons was detected after 24 hours of incubation with STEP-RNP. Top 3 hsgRNAs (5, 7, 16) with G5 STEP-G5 Cas9 variant and hgRNA-7 with V3 and G221 Cas9 variant and V3 Cas9 (n>6 per condition).

**(e)** STEP-RNP editing of UBE3A-ATS did not affect with the expression of *SNORD116* and *SNHG14(IPW)* but increased the expression of SNORD115 and SNRPN that may be by a compensatory mechanism detected by real time RT-PCR.

**(f)** Comparison of immunofluorescence quantification of 3 versions of Cas9 (V3, G5, and G221) synthesized with hsgRNA-7 on the reactivation of UBE3A-GFP showed that G5 significantly better than V3 and G221 Cas9.

#### Extended Data 5

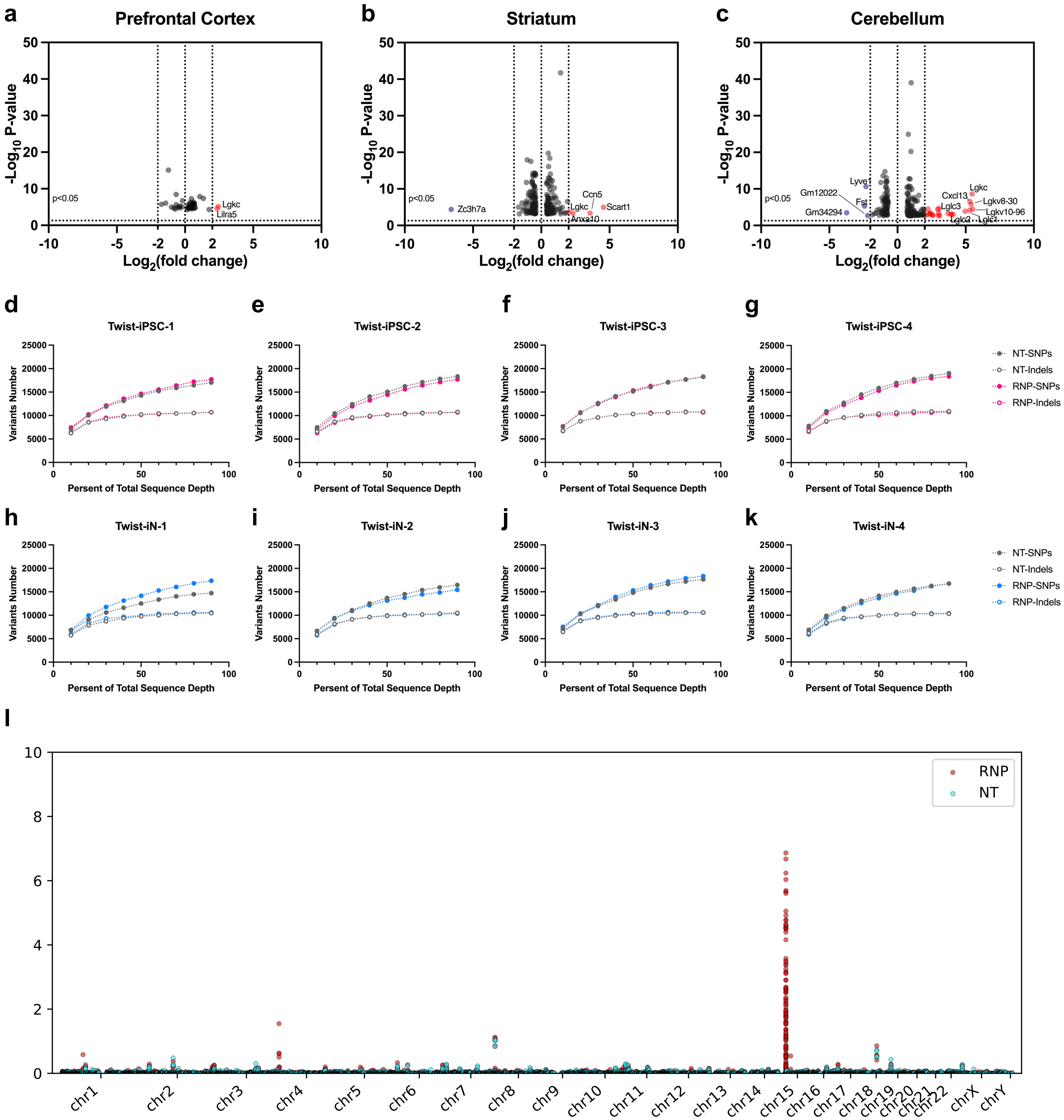

**Extended Data 5**

**Off-target assessment by RNA-seq of mouse brain tissue, CHANGEseq nomination in hIPSCs followed by enrichment hybridization capture in hIPSCs and hIPSCs-induced Neurons.**

**(a-c)** Illumina bulk RNA-seq of the brain tissues from the same mouse of 300X WGS revealed no significant changes in gene expression for *Zfp33b* and *Mroh2a* associated with these off-target events. The reduced or increased expression ($p<0.05$) are revealed for several genes in different brain regions, but no corresponding change at DNA levels are found.

**(d-k)** Saturation analysis of sequencing depth of hybridization capture. Raw variants called from Twist enrichment hybridization panel applied to hIPSCs **(d-g)** and hIPSCs-induced neurons (iN) **(h-k)** from 4 health donors, divided by 10% bins of entire sequence data of each samples. The number of Indels calling saturated at about 30% among all samples in both non-treated (NT) and STEP-RNP treated (RNP) groups, while the SNPs approach to saturation within current sequencing depth too.

**(l)** Manhattan plot of genome-wide on-/off-target evaluation in 4 iN samples. The x-axis represents the chromosomal localization of indels, while the y-axis indicates their frequency. Hybrid capture analysis revealed independent on-target indel frequency of up to 7.062% and an accumulated editing efficiency of up to 53.039% within the chr15:25193313–25439024 (UBE3A-ATS) region.

### Extended Data 6

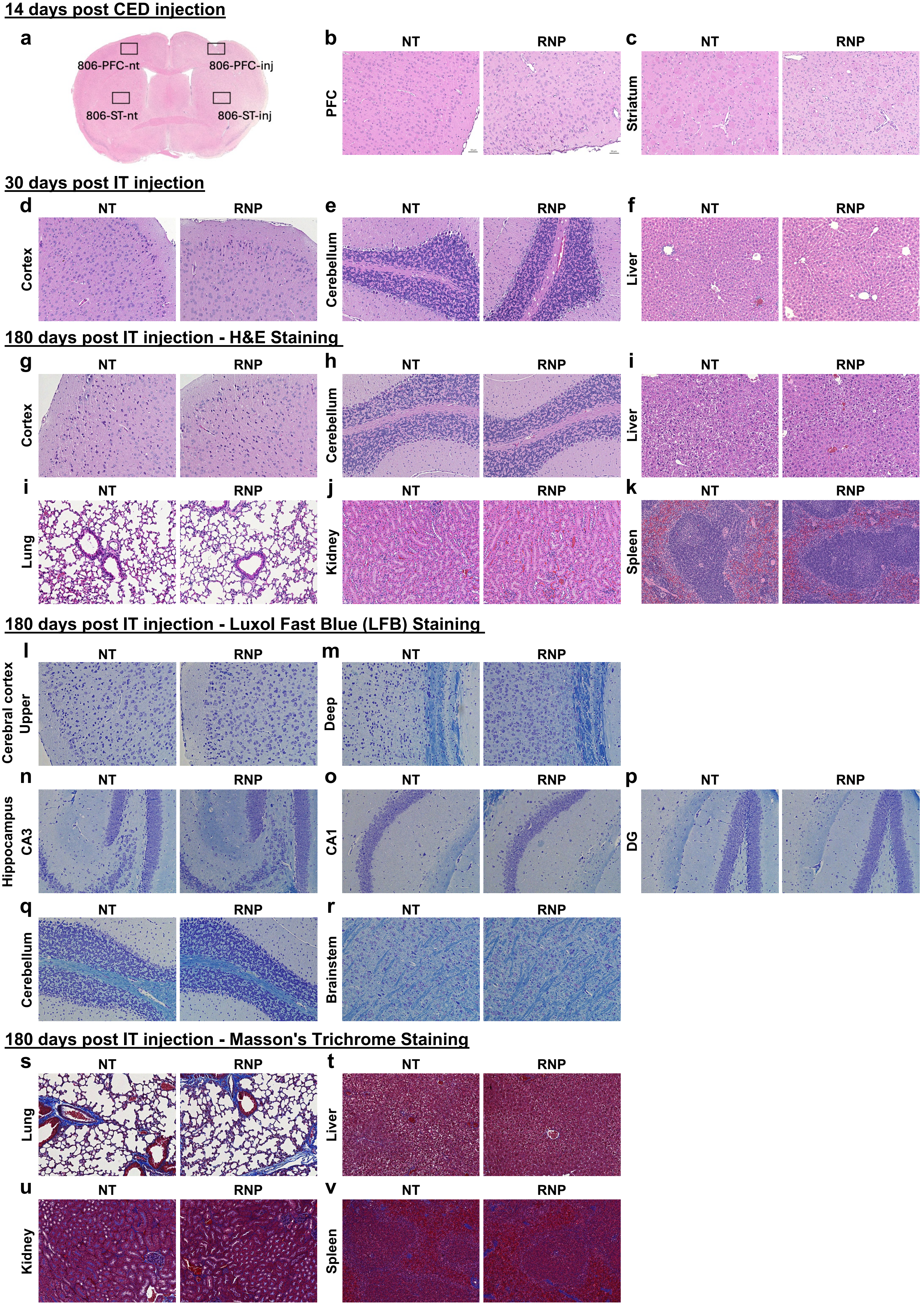

**Extended Data 6**

**Histological analyses of H&E staining (a-k), Luxol Fast Blue staining, and Masson's Trichrome Staining of major organs after administration of STEP-RNP**

**Histological analyses of H&E staining (a-k)**

**(a-c)** Representative image of CED injected Ai9 mouse brains. Samples were collected 2 weeks after injection. Three examples of general histology performed on CED injected mouse brains to assess the toxicity of STEP-RNP. Zoomed images of prefrontal cortex (PFC) and striatum (ST) of injected side on right compared to non-injected side on left.

**(d-f)** Representative image of YFP mice brain (D) and liver (E) 30 days after STEP-RNP administration via IT on P1

**(g-k)** Representative image of cortex, cerebellum, liver, lung , kidney, and spleen of WT mice 180 days after IT administration of STEP-RNP on day1. These straining revealed comparable tissue architecture across all regions, with no observable differences between NT and RNP-treated samples. In the brain, both the cortex and cerebellum exhibit normal cytoarchitecture, with no apparent signs of neuronal loss or structural disruption. The liver shows normal hepatocyte organization, the lung exhibits intact alveolar structures, and the kidney displays well-preserved glomeruli and tubules. The spleen maintains distinct white and red pulp regions without signs of abnormal infiltration. These findings indicate no detectable histopathological changes between the NT and RNP-treated groups. (N=4 RNP treated group and scale bar = 50 um).

**Luxol Fast Blue (LFB) Staining comparison between NT and STEP-RNP Treated mouse brains 180 days after P1 IT administration**

**(l-r) Representative images of** LFB staining of myelinated regions in the cortex, hippocampus (CA1 and CA3), cerebellum, and brainstem in NT and RNP-treated mouse brains 180 days after STEP-RNP administration in WT mice. Myelin staining appears consistent across all regions, with no apparent differences between NT and STEP-RNP-treated samples. The upper and deep layers of the cortex, hippocampal subfields (DG, CA3 and CA1), cerebellum, and brainstem all exhibit comparable staining intensity and structural integrity, indicating no observable demyelination or myelin damage resulted from STEP-RNP administration. (n=4 RNP treated group)

**Masson’s Trichrome staining of lung, liver, kidney, and spleen tissues in NT and STEP-RNP treated mice 180 days after P1 IT administration**

**(s-v)** Masson's Trichrome staining was performed to assess collagen deposition (blue), muscle fibers (red), and cellular structures (purple/dark red) in lung, liver, kidney, and spleen tissues from NT (non-treated) and RNP-treated mice at 18 0 days. In the lung panel, alveolar spaces, bronchi, and blood vessels appear comparable between NT and RNP-treate d groups, with no evident increase in collagen deposition or fibrosis, suggesting that STEP-RNP treatment does not ind uce extracellular matrix remodeling in lung tissue. In the liver panel, hepatic architecture remains preserved across gro ups, with no excessive collagen accumulation in periportal or pericentral areas, indicating the absence of fibrosis. In the kidney panel, renal structures, including glomeruli and tubules, appear intact, with collagen localized primarily to expect ed regions such as blood vessels and interstitial spaces, without excessive deposition in the STEP-RNP-treated group. In the spleen panel, the distinction between white and red pulp is maintained, with minimal collagen deposition restricte d to trabeculae and capsular regions, consistent with normal splenic architecture. Overall, these results indicate that ST EP-RNP treatment does not significantly alter extracellular matrix composition or induce fibrosis in these major organs.

### Extended Data 7

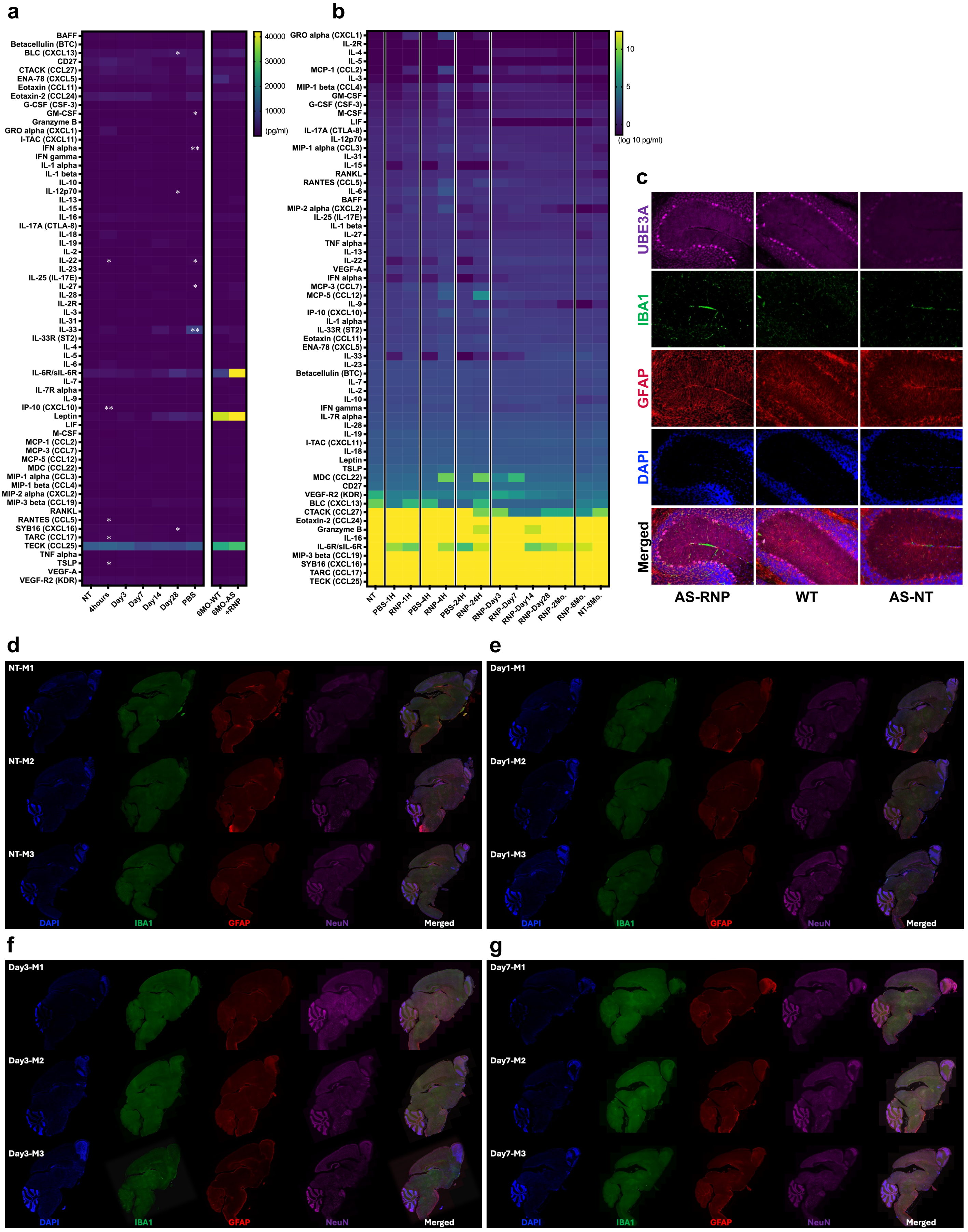

**Extended Data 7**

**Assessment of general toxicity and immunogenicity of STEP-RNP delivery**

**(a)** Immune and inflammatory responses were assessed by protein-based panel in extracted serum of P21 WT mice treated intrathecally IT with STEP-RNP-sgRNA-NC and PBS only or untreated at different time points (4 hours, 3 days, 7 days, 14 days, 28 days and 6 months) .Significant differences were detected for several cytokines (e.g. IL-22, IP-10 and others) at 4 hours following treatment with STEP-RNP-sgRNA-NC and PBS only, compared to no treatment (NT) control. No significant differences were observed at 3, 7, 14 days, but several cytokines were increased on day 28 (NT, n=12; WT 4 hours, 3, 7 days, n=6; WT 14, 28 days, n=8; PBS, n=4). However, no difference between WT and STEP-RNP-AS treated groups at 6 months of age after P21 IT administration (N=6).

**(b)** Immune and inflammatory responses were assessed by protein-based panel in brain tissues of P21 WT mice treated via IT with STEP-RNP-sgRNA-NC and PBS only or untreated after different time period (1h, 4 h, 1 day, 3 days, 7 days, 14 days, 28day, 2 months, and 8 months).

**(c)** Zoomed in view of immunofluorescence staining of IBA1+ microglia and GFAP+ astrocytes the AS mouse brain 8 months after STEP-RNP administration via IT, corresponding to Fig.6s.

**(d)** Immunostainings of IBA1+ microglia and GFAP+ astrocytes at 1 day, 3 day and 7 days after STEP-RNP administration via IT compared to non-treated (NT) mice.
